# Evaluating Methods for Differential Gene Expression And Alternative Splicing Using Internal Synthetic Controls

**DOI:** 10.1101/2020.08.05.238295

**Authors:** Sudeep Mehrotra, Revital Bronstein, Daniel Navarro-Gomez, Ayellet V. Segrè, Eric A. Pierce

## Abstract

High-throughput transcriptome sequencing has become a powerful tool in the study of human diseases. Identification of causal mechanisms may entail analysis of differential gene expression (DGE), differential transcript/isoform expression (DTE) and identification, classification and quantification of alternative splicing (AS) and/or detection of novel AS events. For such a global transcriptome profiling execution of multi-level data analysis methodologies is required. Each level presents its own unique challenges and the questions about their performance remains. In this work we present results from systematic and consistent assessing and comparing a number of widely used methods for detecting DGE, DTE and AS using internal control “spike-in” sequences (Sequins) in RNA-seq data. We demonstrated that inclusion of internal controls in RNA-seq experiments allows accurate determination of lower bounds detection levels, and better assessment of DGE, DTE and AS accuracy and sensitivity. Tools for RNA-seq read alignment and detection of DGE performed reasonably. More efforts are needed to improve specificity and sensitivity of DTE and AS detection. Low expression of isoforms accompanied with sequencing depth does impact sensitivity and specificity of DTE and AS tools.

## Introduction

Whole transcriptom analysis using next generation sequencing (NGS) technology is a powerful tool in the study of human disease [1–4]. This technology generally referred to as RNA sequencing (RNA-Seq) allows the detection of various types of changes in expression levels. These mainly include differential gene expression (DGE), differential transcript/isoform expression (DTE) and identification, classification and quantification of alternative splicing (AS) and/or detection of novel AS events [5], [6–10]. To perform such diverse and global transcriptome profiling and sample comparisons requires a multi-level data analysis and applying multiple methodologies and software programs. Starting with quality control of the sequencing reads and read alignment, through data normalization and statistical calculations of significance, each level of analysis presents its unique challenges. For example, confident estimation of expression levels and splicing events can be challenged by presence of sequencing errors impacting accurate read alignment, biases from library preparation methodology and complexities from exons shared by overlapping genes and their isoforms [7] and [11]. Isoform reconstruction and quantification is also a challenging task with short RNA-Seq reads, in particular for large genes with multiple exons [12]. Multiple computational tools/software programs have been generated, each with their own statistical approach and assumptions. Various studies have evaluated the performance of available tools used in RNA-Seq analytical workflows at varying levels. Some are limited to alignment and/or quantification of feature(gene/isoform) expression or characterization of splicing. A few studies performed a comprehensive and systematic analysis of the RNA-Seq data. They highlight strengths and weaknesses of different alignments, DGE, DTE and AS tools (not an exhaustive list); [7, 13–19]. Efforts such as by the Sequencing Quality Control Consortium (SEQC) data set and the synthetic RNA spike-in controls from the External RNA Control Consortium (ERCC) have extended our understanding of the pros and cons of different sequencing protocols, sequencing centers and platforms and data analysis pipeline [20] and [21]. The ERCC RNA spike-in controls contribute mostly to differential gene expression profiling and junction discovery. However, they do not mimic the complexities of the mammalian genomes and are thus less able to support the evaluation of tools aimed at discerning and quantifying splicing events and isoform quantification.

In this study we have used internal control “spike-in sequences” — sequins —to test the current capabilities and limitations of widely used RNA-Seq analysis tools [22]. The sequins represent full-length spliced mRNA isoforms, modeling both differential gene expression and alternative splicing events [22]. These synthetic RNAs are designed to model complexities associated with human/mouse transcriptome covering a wide range of gene and transcript isoform expression levels. We added sequins to five (5) different tissues; retinal pigment epithelium (RPE), retina, brain, muscle and kidney from a model organism, mouse, with consistent and realistic sample replicates. We evaluated the performance of the following tools for RNA-seq read alignment – STAR [23] and HISAT2 [24], gene expression quantification – featureCount [25] and HTSeqCount [26], differential gene expression – DESeq2 [27] and edgeR [28], isoform quantification: RSEM [29] and Kallisto [30], differential transcript/isoform expressions – EBSeq [31] and Sleuth [17], and alternative splicing detection – JunctionSeq [32] and MAJIQ [33].

We demonstrate that inclusion of internal controls in RNA-seq experiments allows accurate determination of detection levels, and better assessment of DGE, DTE and AS accuracy. These analyses show that gene expression profiling and DGE tools were found to be robust. The isoform expression profiling tools were robust, but not as tolerant to lower depth of coverage (DOC) as the DGE tools. We recommend high, for confident prediction of differential isoform expression. Even with high DOC, for AS event detection, we found the current tools lacking high sensitivity. This was compounded by lower depth of coverage. More efforts are needed to improve specificity and sensitivity of DTE and AS detection.

## Materials and Methods

### Data Sets: Isolation & Sequencing

The RNA-Seq data used for the analyses described were generated from tissue of mice with targeted disruption of the *Prpf31* gene [34] and [35]. Total RNA was extracted from retinal pigment epithelium (RPE), retina, brain, muscle and kidneys of 5 *Prpf31*^+/−^ and 4 wild-type littermate male mice (C57BL/6J genetic background). Immediately following euthanization with Co_2_, brain muscle and kidney were dissected, washed in PBS and flash frozen in liquid nitrogen. Tissues were lysed using Geno/Grinder (SPEXSamplePrep Metuchen, NJ, USA). Eyes were enucleated and RPE and retinas were isolated as previously described by Fernandez-Godino et al. [36]. Isolated retinas and RPE monolayers were collected into buffer RLT and placed into −80C. Total RNA was extracted from brain, muscle and kidney using RNeasy maxi kit and from retinas and RPE using RNeasy mini and micro kits respectively (Qiagen, Hilden, Germany) following the manufacturer instructions. RNA was quantified using Qubit (ThermoFisher scientific Waltham, MA, USA) and quality was assessed using a Bioanalyzer (Agilent Santa Clara, CA, USA). RIN numbers were *>*7 for all the samples.

### Data Sets: Library Preparation & Sequencing

For each retina, brain, muscle and kidney sample, 1*µ*g of total RNA was spiked with 1.2ng sequins (v2) controls. For RPE samples 100ng total RNA was spiked with 0.12ng sequin controls. Sequins “MixA” was added to all the wild type (WT) samples and “MixB” to mutant type (MUT) samples. A stranded, paired-end (PE) TruSeq stranded total RNA (Illumina San Diego, C, USA) library preparation was performed. Ribosomal RNA was removed with the Ribo-Zero Human/Mouse/Rat kit. Samples were multiplexed across five different flow cells and were consecutively sequenced. The sequencing was carried out on Illumina HiSeq 2500 Sequencer for 101 cycles at the Ocular Genomic Institute (OGI), Boston, MA (USA).

### Preprocessing and Sample Quality Control

Each section of the analysis pipeline was run on a local High Performance Cluster (HPC) with SunGird Engine, 12 compute nodes each with 16 core processor (2X8) and 128 GB memory and the Ocular Genomic Institute (OGI), Boston, MA (USA). First we performed preprocessing of the data set. The quality control/quality assurance (QC/QA) of the data set at lane and samples level. Our aim was to identify any lane level or sample biases in pre-processing of the sequenced reads.

The quality of the sequenced reads was evaluated using FastQC and further collated and summarized using MultiQC [37], [38]. Post merging, an in-house Perl script was used to check and filter out reads with presence of adapter and ambiguous character, ‘N’. The Bowtie2 aligner was used to identify ribosomal contamination [39]. Reads aligned to rRNA reference sequences were dropped from all downstream analysis using SAMtools [40]. The mouse (*Mus musculus*) reference sequence was downloaded from Ensembl and annotation files from GENCODE. Supplementary Table S1 lists all the reference databases along with the versions that were used in the analysis. Additionally, Supplementary Table S2 shows all the analysis tools along with versions that were used. Post alignment, featureCount was used to generate gene expression matrix [25]. To identify possible mixing/mismatch, the normalized read counts were used for clustering of all the samples using Spearman correlation. In summary no lane level or sample level biases were observed. No samples mixing was detected (data not shown).

### Comparative Analysis

In the paper performance of various tools used in RNA-Seq work flow are analyzed. All possible combinations of aligners, gene expression profiles, differential genes and isoform expression tools are analyzed. Following sections expands on the tools used in the analysis.

#### Overview of Sequins

A suite of synthetic RNA isoforms, termed “sequins” (sequencing spike-ins) representing full-length spliced mRNA isoforms, which are entirely artificial sequence and bore no homology to natural reference genomes (human/mouse) have been designed by Hardwick et al. [22]. They align to gene loci encoded on an artificial *in silico* (IS) chromosome. The sequins can be concatenated with the genome of interest (such as human and mouse) and coindexed for alignment. In total, there are 78 artificial gene loci encoding 164 alternative isoforms comprising 869 unique exons and 754 unique introns. Genes ranged from single-exon to large multi-exon loci, with individual isoforms ranging in size from *∼*280 bp up to *∼*7 kb and comprising up to 36 exons. For more information please refer to [22].

#### Overview of Analysis Tools: Aligners

The aligners are central and primary in any RNA-Seq analysis. Two different state-of-the-art splice site aware aligners, STAR and HISAT2 were used for alignment [23] and [24]. The alignment was performed using default settings. The STAR aligner was used to align in two-pass mode within the sample and across replicates for each sample set. Respective indexes were created for the mouse reference as per the manual. During preprocessing and QC/QA the reads were aligned using STAR.

#### Overview of Analysis Tools: Gene Expression

Identification of differentially expressed genes between case and control is vital to many RNA-Seq studies. The first step in this process is creation of a count based profile of expressed genes. A profile refers to genes its read counts across all the samples and conditions. In this study featureCount (from the subRead package, and sometimes referred to as “subRead” in the paper) and HTSeqCount (htseq-count) were used to build gene expression matrices [26]. Mostly, default settings were used for HTSeqCount, calibrating on parameters for reverse-stranded and PE reads as required. The following non default settings for featureCount were applied: (i) reads must be paired (ii) both the pairs must be mapped (iii) use only uniquely mapped reads (iv) multi-mapped reads are not counted and finally (v) chimeric reads are not counted.

We show Pearson’s correlation of determination (R^2^) via Anaquin or using R. In come cases, marked appropriately, for correlation and other plots the transcripts per million (TPM) for each sample or mean TPM values across the appropriate genotypic replicates were used. All the correlation plots were made using “Corrplot” [41].

#### Overview of Analysis Tools: Differential Gene Expression

DESeq2 and edgeR are R/Bioconductor packages and were used for differential gene expression (DGE) analysis [27]. Both the tools were used with default parameters. Both the tools apply a negative binomial generalized log-linear model to identify differential expression between experimental conditions.

#### Overview of Analysis Tools: Isoform Expression

As with genes, identification of expressed transcripts requires creation of expression profiles. For isoform analysis, RSEM and Kallisto were used [29] and [30]. For RSEM we used alignments from Bowtie2 (default for RSEM), STAR and HISAT2 aligners as input to create isoform expression profiles. Kallisto intrinsically uses a “pseudo-alignment” approach distinct from splice site aware aligner (STAR/HISAT2) for detection of isoform expression. In all the tools, calibrations to the parameters were made for reverse-stranded and PE reads. As with genes, for correlation and other plots the read counts were first converted to TPM and mean values were used.

#### Overview of Analysis Tools: Differential Transcript/Isoform Expression

For differential transcript/isoform expressions, EBSeq and Sleuth were used. RSEMs expression profile matrix were used as input for EBSeq while Kallsito’s output were input to Sleuth for DTE. EBSeq applies the Bayesian empirical approach for DTE. Sleuth relies on variance decomposition to better capture the biological differences in transcript, applying shrinkage to stabilize variance across samples replicates [17] for quantification. Both EBSeq and Sleuth were used with default parameters. Note that Sleuth use likely-ratio-test (LRT) in default mode which does not produce LFC values. It allows “Wald” testing as well. We followed the same methodology as described by Pimentel et al [17] for data preparation for LFC and receiver operating characteristic (ROC) for comparison.

#### Overview of Analysis Tools: Alternative Splicing

We used count-based methods that include both exon-based and event-based approaches. In exon-based methods, read counts are assigned to different features, such as exons or junctions. JunctionSeq is an R/Bioconductor package that applies such an approach and is capable of detecting novel exon junctions. “Differential usage” (DU) is an observed phenomenon in which individual exons or splice junctions display expression that is different with the overall expression of the gene [32]. Hence, the limitation of JunctionSeq is it that it does not make any inference on the type of the splicing event in classical terms such as; Exon Skipping(ES), Intron Retention (IR), Mutually Exclusive Exons (MXE), Alternative 5′ Splice Sites (A5SS), Alternative 3′ Splice Sites(A3SS), Alternate first exon (AF) and Alternate last (AL) exon, which are easier to interpret in the biological and universal manner. In an event-based method, splicing events are quantified by calculating the percentage spliced in (PSI) values for each event by measuring the differences in the fraction of junction reads, followed by calculation of difference in PSI (dPSI) between experimental conditions. MAJIQ (Modeling Alternative Junction Inclusion Quantification) [33] uses a local splicing variations (LSVs) approach to detect and quantify RNA splicing. The event detection terms utilized by MAJIQ are restricted to ES, IR, A5SS and A5SS. MAJIQ uses a Bayesian approach and reports posterior probability per LSV as confidence values. Another tool, rMATS [42], is also an event-based method for splice detection and uses a likelihood ratio test to calculate the *P* value and examines whether the between-group differences of mean PSI exceed a given, user-defined threshold. Non-default settings for minimum of 5 reads for junction detection and 10 reads for the calculation of dPSI. Additionally, calibrations to the parameters were made for reverse-stranded and PE reads. Using sequins sensitivity and specificity of every parameter can be tested. However such an analysis is beyond the scope of this paper. We focused on the systematic reporting of the event across and the tissues and the location rather than on the type of AS events.

#### Downsampling

To explore impacts of lower depth coverage (DOC) on DGE, isoform expression and AS events, we titrated the original unique alignments into random 150, 100 and 50 million set. The downsampling was performed on RPE, retina and brain samples. For each sample regardless of the genotype we first determined the samples that can undergo downsampling. Not all samples had enough depth of coverage to meet the criterion to downsample. In such instances, the entire sample was used. To ensure proper sequins ratio, for each of these samples, observed (post downsampled) total sequins counts was extracted and compared with the original set. We changed the “seed” during random selection for downsampling.

## Results

The RNA-seq data used for the analyses described below were generated from total RNA extracted from RPE, retina, brain, muscle and kidney of mice with targeted disruption of the *Prpf31* gene and littermate controls. Mutations in the *Prpf31* gene cause inherited retinal degeneration [35]. In *Prpf31*^+/−^ mice this manifests as cell autonomous defects in retinal pigment epithelial (RPE) function [43]. Two different concentration mixes of sequin spike-in controls were included in the RNA samples isolated from 5 tissues of the mutant and control mice with 5 and 4 replicates respectively (Methods).

We used these RNA-seq data to comprehensively evaluate landscape of various bioinformatics tools used in RNA-Seq experiment from read alignment to gene expression quantification and their differential analysis, isoform expression quantification and their differential analysis and alternative splicing detection. The strategies and various analysis tools used for comparative analysis in this paper are illustrated in Figure 1.

**Figure (1).**
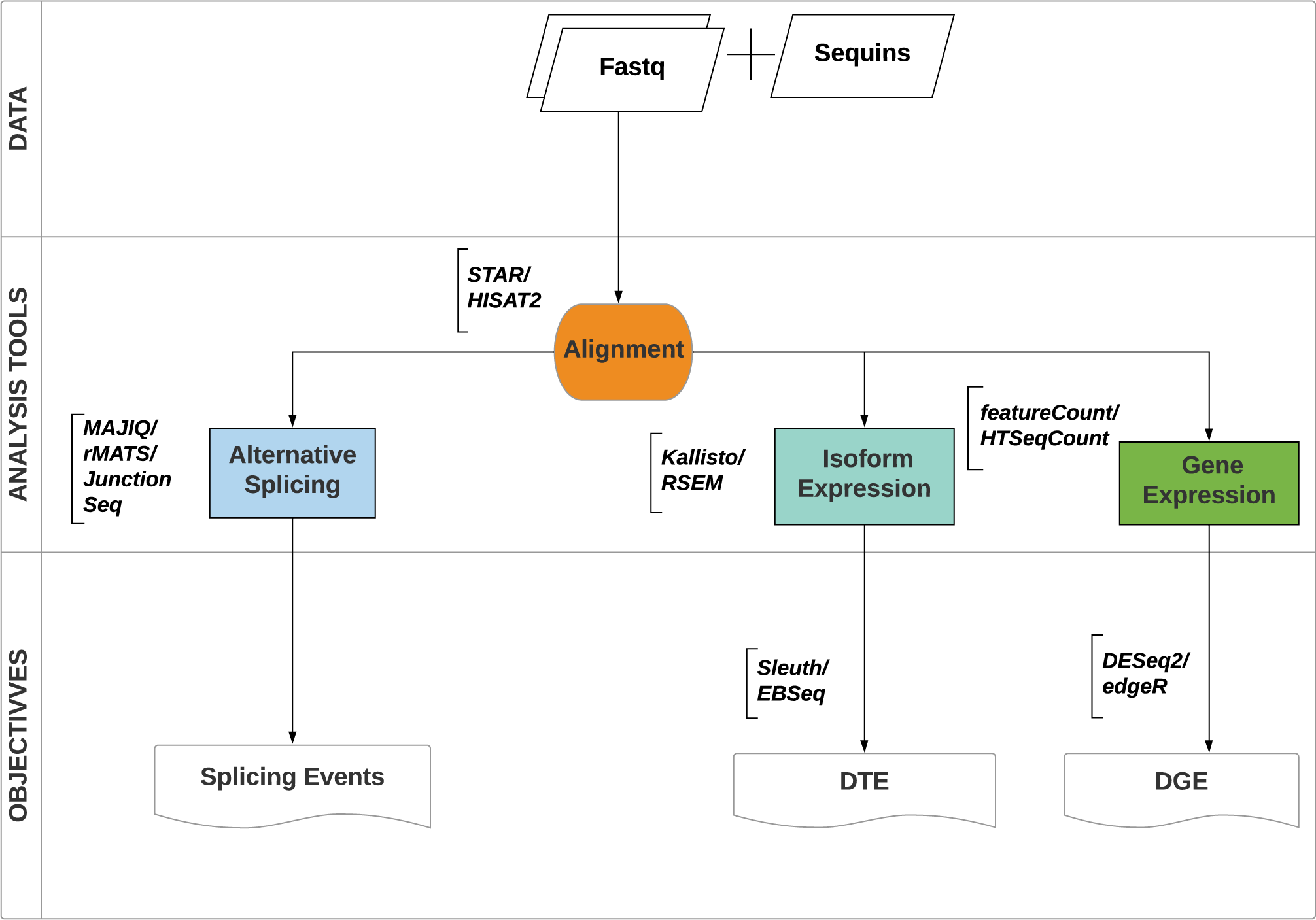
Experimental design flow chart. Top: The RNA-Seq data in the form of fastq files were used as input to alignment tools. Post alignment, different combination of software programs were used for gene expression, isoform expression profiles were created including detection of splicing events, with objective to perform different gene expression (DGE), differential transcript expression (DTE) and differential splicing events.

### Sequencing Quality

Since the samples were multiplexed and were consecutively sequenced across 5 flow cells, we tested for any lane biases so as to avoid additive biases that can occur following merging of the reads. During QC/QA aspects such as: total raw reads, sequence quality, total number of bases, GC content, the presence of adaptors and overrepresented k-mers in order to detect sequencing errors, PCR artifacts or contaminations are paid particular attention. Supplementary Figure S1 shows total raw reads including a comparative with high quality (HQ) read set post QC. Here, reads and bases that underwent robust QC process are referred to as HQ. Post QC, 98%–99% reads were retained across all the sample set for all the tissues. Furthermore, Supplementary Table S3, Table S4,Table S5,Table S6 and Table S7 shows the mean, first and third quartile values of each sequencing run. Overall no lane level biases were observed. Supplementary Figure S2 shows mean per cycle per base quality. The data used for subsequent analyses contained high base quality data.

### Sequins In Samples

Sequins are provided as two mixes which differ in gene expression, transcript isoform usage and levels [22]). In this study, mixA was added to all the wild-types sample replicates and mixB to all the mutant sample replicates for all the mouse tissue types. Table 1 shows the final average ratios of sequins across the mouse tissues and genotypes.

**Table (1).** Detected average sequins ratio in case and control samples. The detected ratios are shown for each mouse tissue. With exception of brain tissue where a disproportionate of the two sequins mixtures were observed. About 35% of the mixA reads for mutants and 65% of mixB reads for the wild types were observed. In all the other tissues, equivalence of the two mixes was observed. IS refers to artificial sequins *in-silico* (IS) chromosome.

| Tissue/Ratio | RPE | Retina | Brain | Kidney | Muscle |
| --- | --- | --- | --- | --- | --- |
| %Avg Ratio<br>MUT:WT | 3.4:3.9 | 4.2:4.5 | 2.7:5.0 | 3.9:4.5 | 3.8:4.2 |

Two sequin genes from the original set were identified to be inconsistent (confirmed by personal communication) and were removed from the set (gene model file). Supplementary Table S8 shows sequins for gene expression and DGE. All the 76 true positive (TP) genes, for upregulation, downregulation and uniform (no change) are shown. The table shows expression values in original attomoles/*µ*l units. Similarly, Supplementary Table S9 shows the TP set of sequins for DTE. Concentration shown are the same as genes. Finally, Supplementary Table S10 shows TP sequins for AS events. In total we have 28 TP events. However, regardless of the use of aligner, 6 sequins (colored in the table), were not well covered, we did not penalized sequin genes for the lack of coverage. We revised the TP counts for AS events to 22. Supplementary Figure S3 and Figure S4 shows the coverage plot for one such sequin gene, R1_62. This is not among the lowest concentrations (8th highest concentration). For brevity of the 6, one(1) such gene with low coverage is shown. The remaining low coverage genes are reflective of this (R1_62) sequin gene.

### Consistencies of the Aligners

Two aligners, STAR and HISAT2 were used for the alignment for all the five tissue types. Figure 2 shows the unique alignment rate from the two aligners using two representative samples; RPE and retina. The alignments were performed to the mouse genome and to the sequins artificial chromosome (IS).

**Figure (2).**
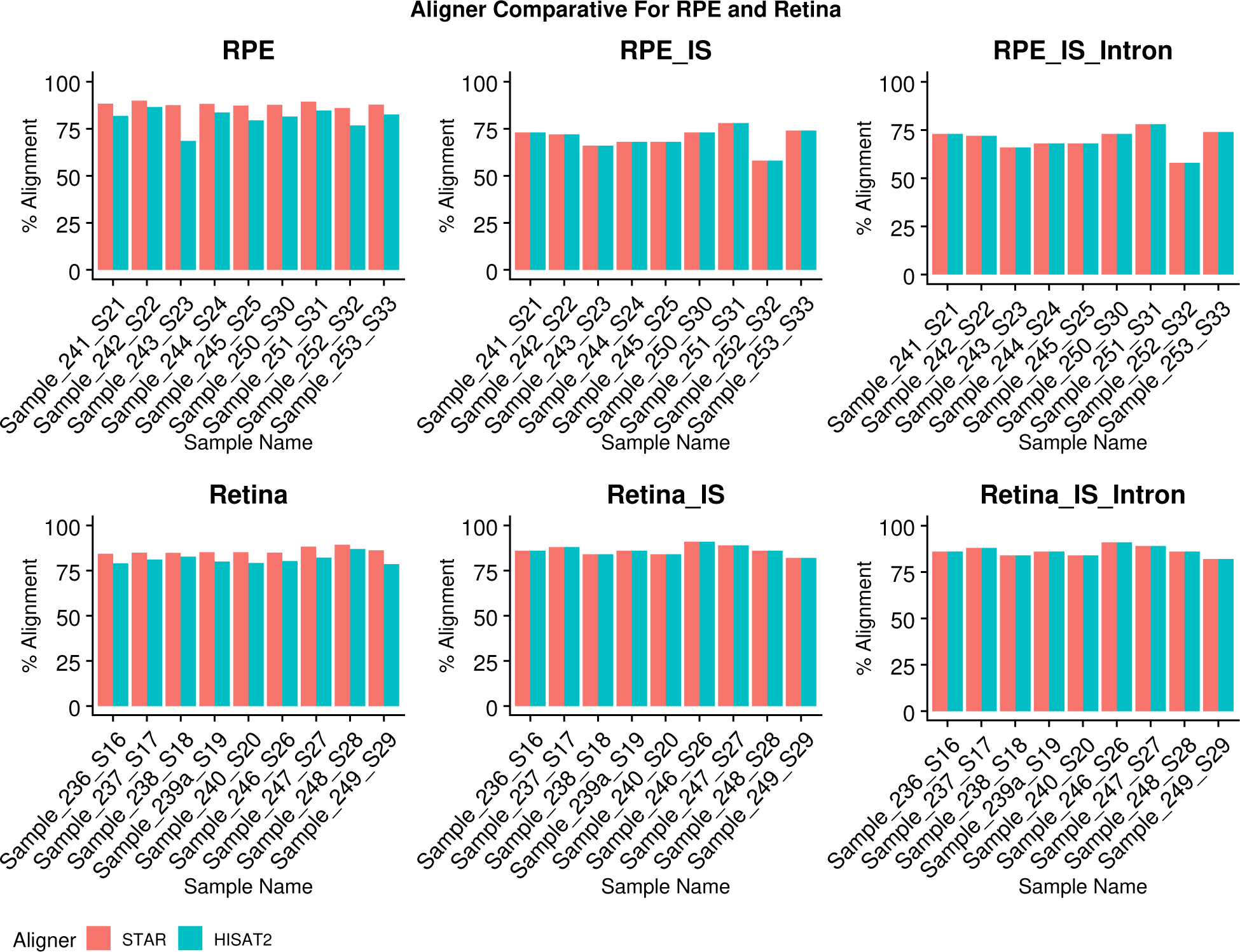
Comparative alignment rate for all the replicates. The figure shows genomic unique alignment, alignment to the sequins artificial chromosome (IS) and intron sensitivity (IS), as reported by STAR and HISAT2 for all the RPE and retina mouse tissues. Overall the alignment rates were indistinguishable. Assigned unique sample identifier assigned to each sample is shown in the x-axis and percentage alignment rate on y-axis. From left to right, first 5 are mutants followed by 4 wild type samples. Supplementary Figure S5 shows genomic alignment, Figure S6 shows alignment rate to *IS* and Figure S7 shows intron sensitivity from the two aligners for all the five mouse tissues.

Since the intron boundaries for the sequins are known, using Anaquin, intron sensitivity was also calculated. Here, sensitivity indicates the fraction of annotated regions covered by alignments of the reads as reported by the aligners. Supplementary Figure S5 shows genomic alignment, Figure S6 shows alignment rate to *IS* and Figure S7 shows intron sensitivity from the two aligners for all the five mouse tissues. Proportionate alignment percentage was observed from both the aligners for all the tissues irrespective of the genotypes and the references.

In the gene model for the spike-in sequences, total sequin gene counts and their relative expression values are known. We used this information to determine following quantitative and qualitative values: limit of detection (LOD) and limit of quantification (LOQ), correlation of determination of the expected versus observed expression values and finally, correlation of determination of log-fold-changes (LFC) and their proportions across the five different tissue types. Over the next few sections aforementioned aspects are explored and results are presented.

### Concentration Levels and High Gene Expression Correlation

The two performance characteristics related to analyte stability, LOD and LOQ were analyzed for all the five tissues sample replicate sets using Anaquin (see [22] and [44] for more details). Figure 3 shows correlation and slope for mouse RPE. This tissue had the lowest depth of coverage amongst all the tissue types and hence was selected as a representative sample set for LOD and LOQ reports. No LOD and LOQ were reported for all the remaining mouse tissues.

**Figure (3).**
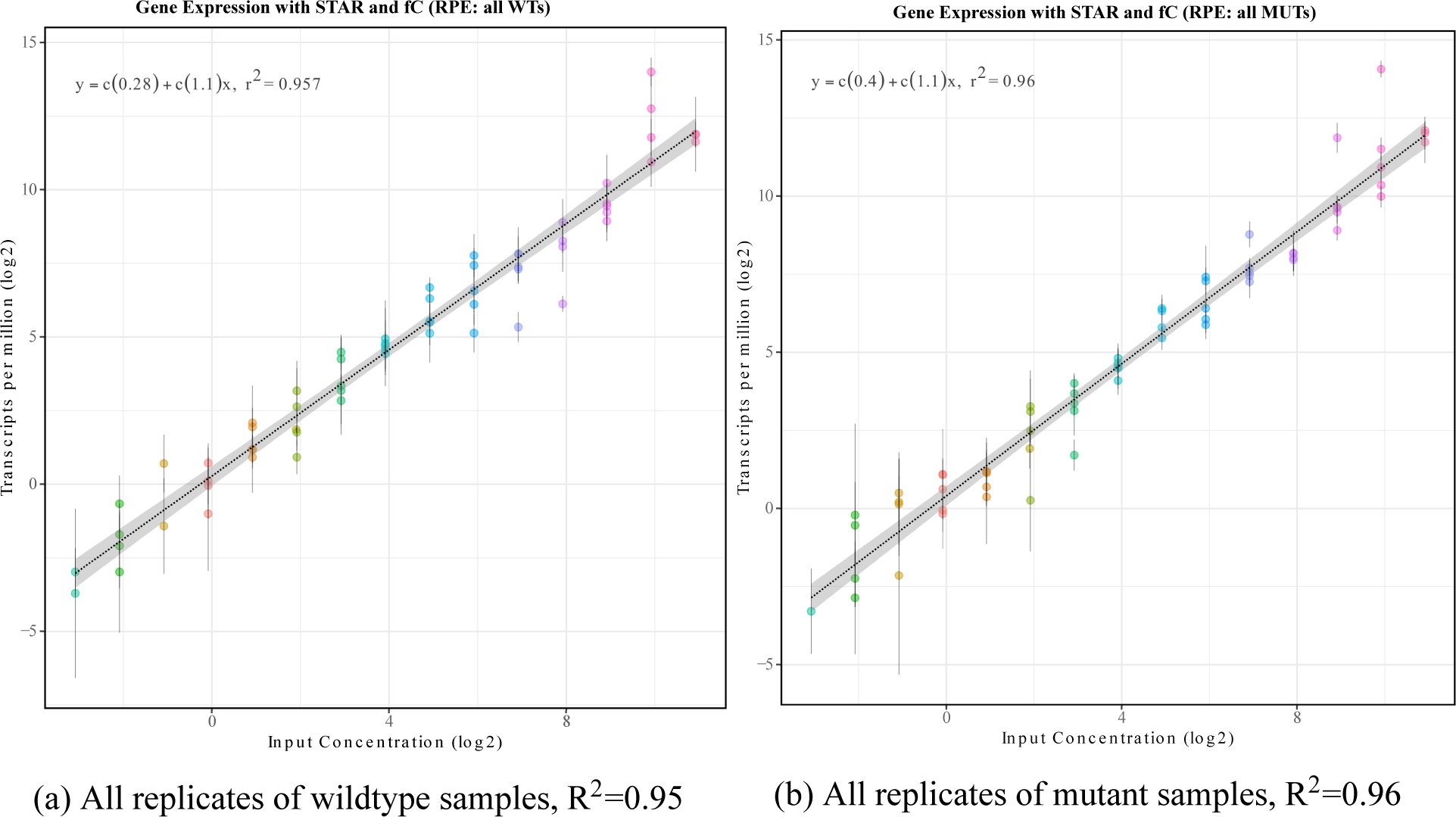
Correlation plot for comparative analysis across all the mouse wild type tissues (a) and mutant tissues (b). Slope, correlation, LOD and LOQ determination using STAR is shown. Known concentrations are on the x-axis and observed concentrations in TPMs is shown on the y-axis. Both the axis are log2 scaled. The vertical bar shows for each sequin genes shows the spread of the concentration values. No LOD or LOQ was detected for this sample set. Few sequin genes were either under or over represented from the slope. fC refers to featureCount.

Next, we calculated R^2^ to determine how well, post alignment, read counting tools; featureCount and htseq-count are able to capture the features and their expression values across the sample replicates. Indeed, all combinations of aligners and read counting tools provided the same high correlation across all samples. Figure 4 and Supplementary Figure S8 show correlation values across the WT and MUT replicates respectively using all possible combinations of the aligners and read counting tools.

**Figure (4).**
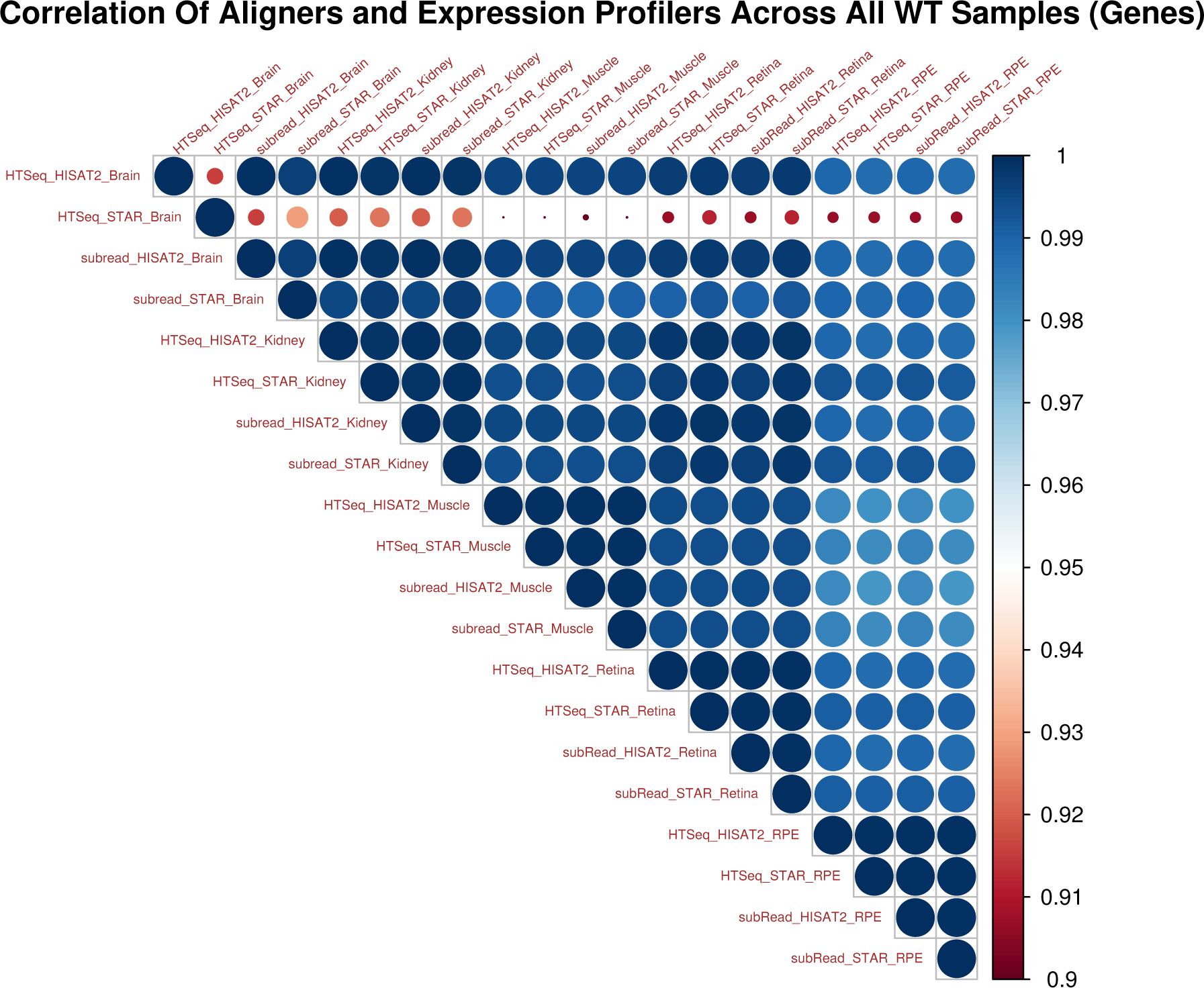
Correlation plot for comparative analysis across all the wild type tissues. An upper triangular heatmap shows high correlation across different combinations of the aligners and genes read counting tools across the five mouse tissues and the replicates. The average TPM values across the replicates were used to calculate the Pearson correlation. The combinations of tools used are shown in the format of: Gene Expression Profile_Aligner_Tissue. The brain samples were impacted by disproportionate sequins ratios.

### High Differential Gene Expression Correlation and Proportion

Our next objective was to analyze differential gene expression across all the tissue types and case and controls set. For these analyses, we computed the correlation values for two different DGE tools, edgeR and DESeq2. Sequin genes are added in a known concentration across the tissues, hence we can examine the correlation of gene expression (LFC) across different tissues. Figure 5 shows an upper triangular computed correlation of determination in the form of a heatmap.

**Figure (5).**
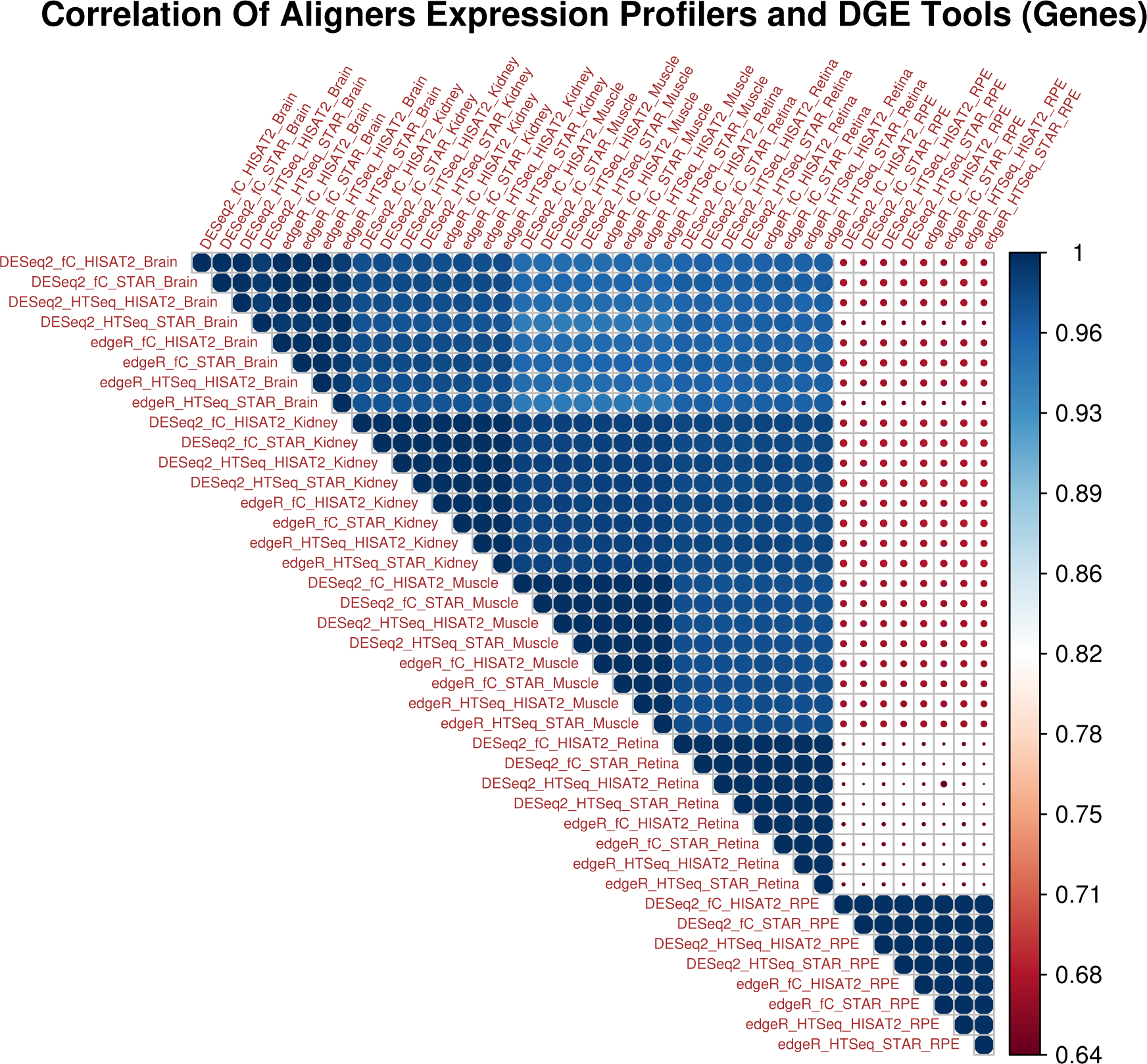
Correlation plot for comparative analysis using two aligners, two read counting tools and two DGE tools across all the mouse tissues. An upper triangular heatmap is generated showing Pearson correlation of determination values using average TPM expression values across the five mouse tissues in the order; brain, kidney, muscle, retina and RPE. The combinations of tools used are shown in the format of: DGE_Gene Expression Profile_Aligner_Tissue, for example, DESeq2_fC_HISAT2_Brain. High correlation of determination values were observed across different tissues types and combinations of different tools with two exception. Correlation of RPE and brain showed the lowest correlation of determination values with other samples. Lower coverage impacted the RPE samples and unequal sequin ratio impacted the brain sample set.

The figure shows (R^2^) values for all possible combinations for the two aligners, two feature read count tools, and two differential gene expression analysis tools across all the five mouse tissues. Overall high correlation of determination values were observed regardless of tissues and tools used. The RPE and the brain samples showed lower values compared to other tissues. For RPE this is likely due to reduced depth of coverage and the brain samples were impacted by disproportionate sequins ratios.

Next, we investigated proportion and directionality of LFC values across all the mouse tissues. Intuition would suggest that for an artificial set of genes, the LFC values and directionality should be proportionate and unidirectional regardless of the mouse tissue type if the depth of coverage is comparable and the ratios of the artificial gene sets were added equally. Figure 6 shows proportions and directionality of sequin genes across all the mouse five tissues. The values shown are sorted based on actual LFC values. Some inconsistencies in the directionality and LFC proportions were observed.

**Figure (6).**
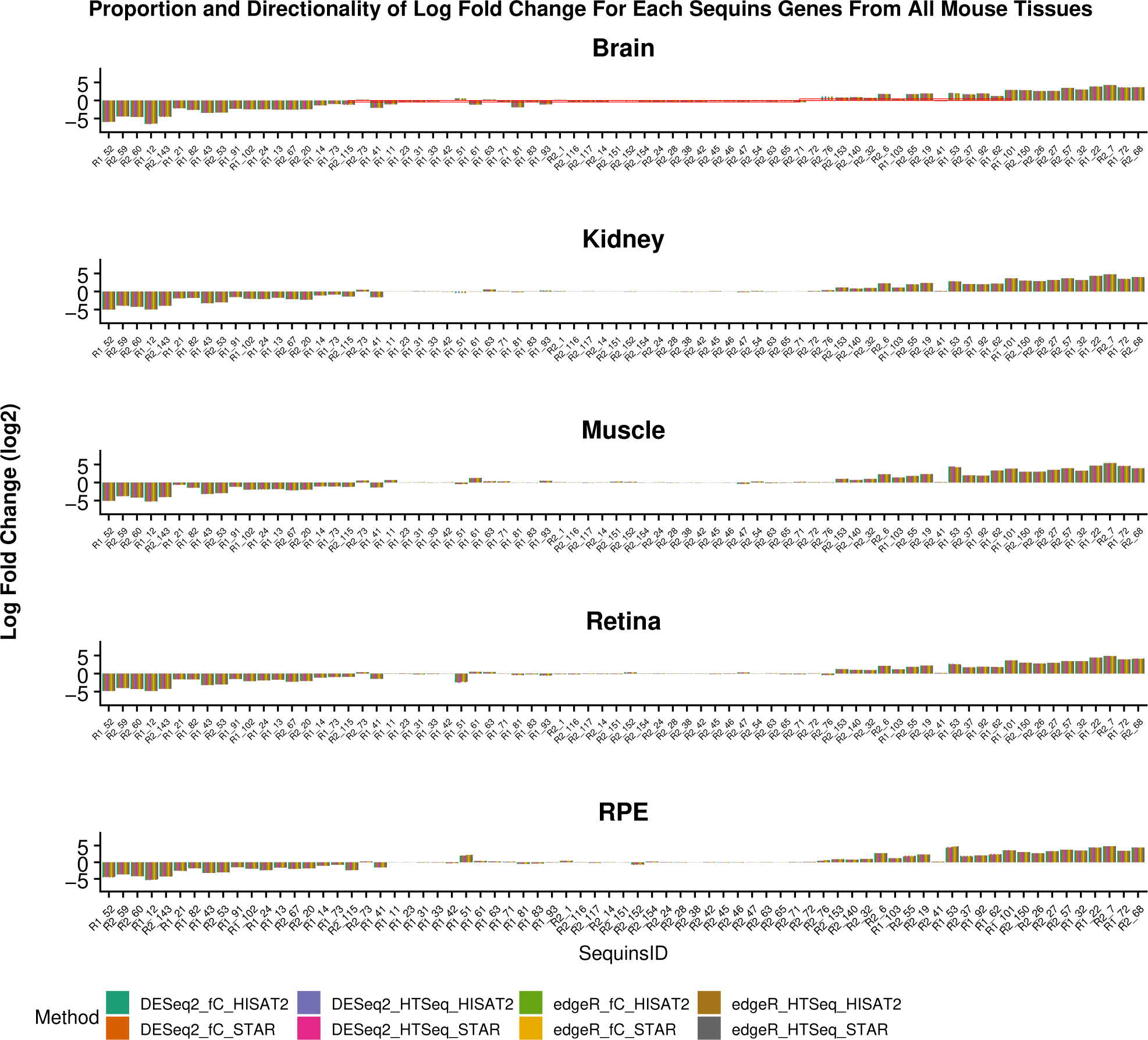
Comparative proportion of log-fold-changes for the sequin genes across all the mouse tissues using different aligners, read counting and DGE tools. In the x-axis all the sequin genes are shown and LFC values (log2) is shown on the y-axis. The values shown are sorted based on actual LFC values. In the top panel, the brain sample highlights three sections (red rectangle boxes) that show some difference in the proportions and in some cases directionality across all the tissues. In most cases, discrepancies in the proportionality and directionality was impacted for sequin genes that should have no (zero) LFC values. In such cases, some marginally over or under LFC values are reported (at most by 1 unit). For example R1_51 was reported 2 units higher in RPE and the same was 2 unit lower in retina. Actual LFC is 0. R1_21 should show about −3 LFC, however in muscle it is barely reaching ~−1. Sequin R1_51 was not reported by some combinations of the tools in the kidney and brain samples. The brain sample set shows the most inconsistent results compared to the rest of the mouse tissues. fC refers to featureCount from the subRead package, HTSeq refers to HTSeq-Count.

In the brain samples, there were set of sequin genes that showed disproportions and changes in directionality. This was mostly observed for sequin genes with ground truth of LFC 0 (zero). Such genes were over or under represented (upto 2 units in either direction). Three sequins were not reported. In the kidney dataset, we observed loss of 2 sequins.

The underlying expected fold changes between the case and control sample sets is known, we can gauge the sensitivity and specificity of various combinations of the tools using receiver operating characteristics (ROC). Figure 7 shows the ROC curve for all the mouse tissues with different combinations of all the analysis tools.

**Figure (7).**
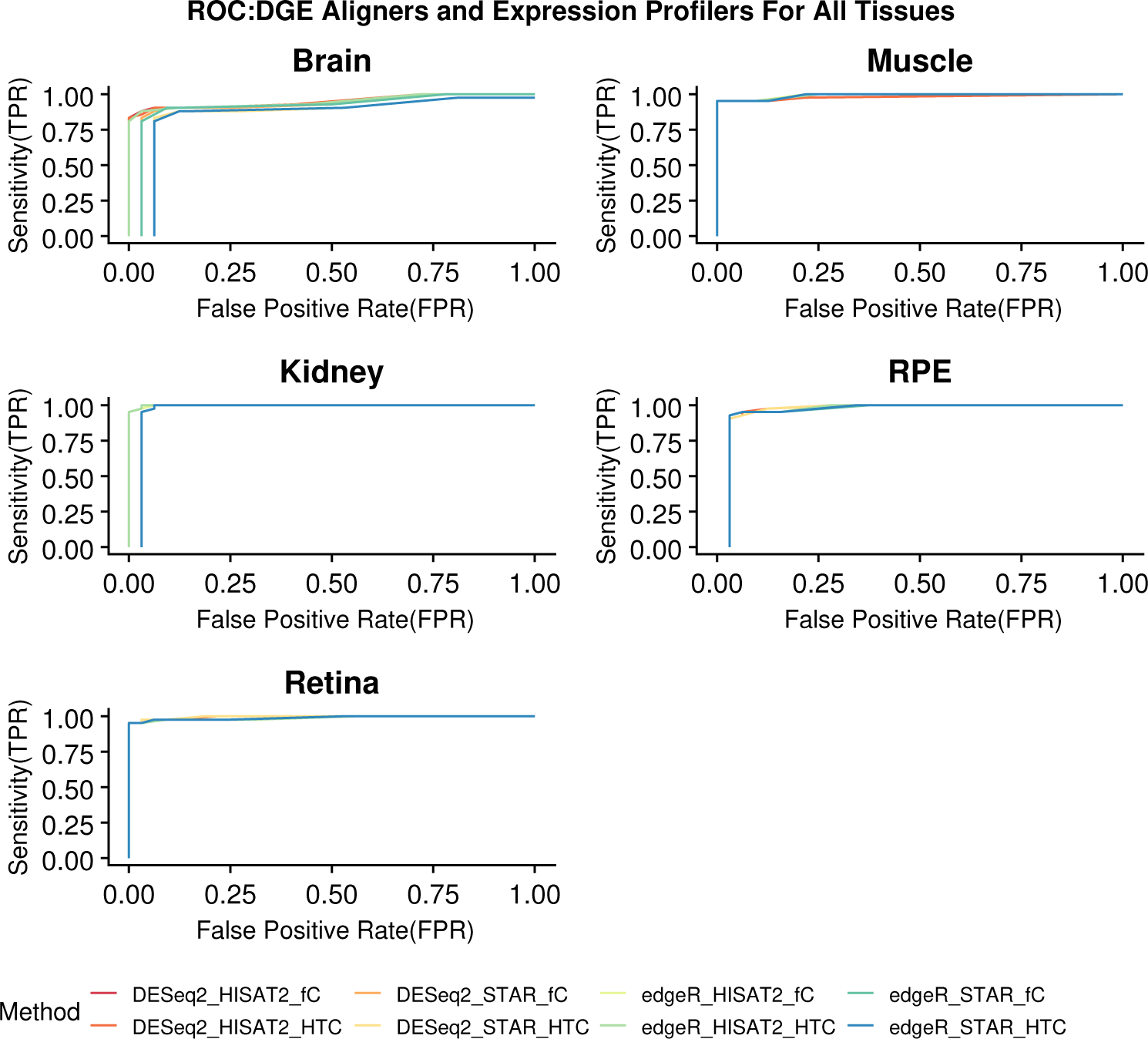
ROC (receiver operating characteristic) curves for Sequin genes for diagnostic performance in measuring fold change. Each plot reflects the performance of different combinations of the aligners, feature read counting and DGE tools. The brain sample set shows an exception compared to the rest of the tissues. The visibility of all the combinations is impaired by the fact that in many cases the results were identical and hence overlapping. fC refers to featureCount from the subRead package, HTC refers to HTSeq-Count.

Each plot in the figure for a given tissue shows the performance of different combinations of the tools used for the alignments, gene expression values and DGE. All show high accuracy with the different combinations aligners, feature read counting and DGE tools tested. The brain sample set was an exception compared to the rest of the samples. The shift in the curves for certain combinations of the tools was due to missed reporting of the sequin genes. The same was observed with the kidney samples. All the other combinations of the tools showed a great balance between sensitivity and specificity. We further investigated the brain and kidney samples to better identify combinations of the tools with drop in performance was observed (Figure S9). Disproportionate ratio of the two sequins mixtures impacted the sensitivity for the brain sample. Loss of 2 sequins in the kidney sample caused the shift ROC.

### High Isoform Expression Correlation

For isoform expression analysis we used different combinations of alignment tools along with an orthogonal approach. We used RSEM with BowTie2 and RSEM with the STAR aligner as a standard approach for detecting isoform expression, and Kallisto’s unique pseudo-alignment approach as an orthogonal approach. We performed multilevel analysis to gauge performance of analyte stability, LOD and LOQ along with correlation of expected vs observed for the isoforms. Figure 8 shows the correlation plot for the mouse RPE WT samples using RSEM and Kallisto. No LOD and LOQ was reported for any of the mouse tissues via any combination of the tool. Some isoforms showed greater expression variability, but remained non specific to any particular tissue types. Figure 9 shows an upper triangular computed correlation of determination in the form of a heatmap for all the wildtype dataset. Simialr observations were made for the mutant sample set (Supplementary Figure S10). The figure shows (R^2^) values for all possible combinations for the two aligners (STAR and BowTie2), “feature” read count tool (RSEM), and Kallisto across all the five mouse tissues. Overall high correlation of determination values were observed regardless of tissues and tools used.

**Figure (8).**
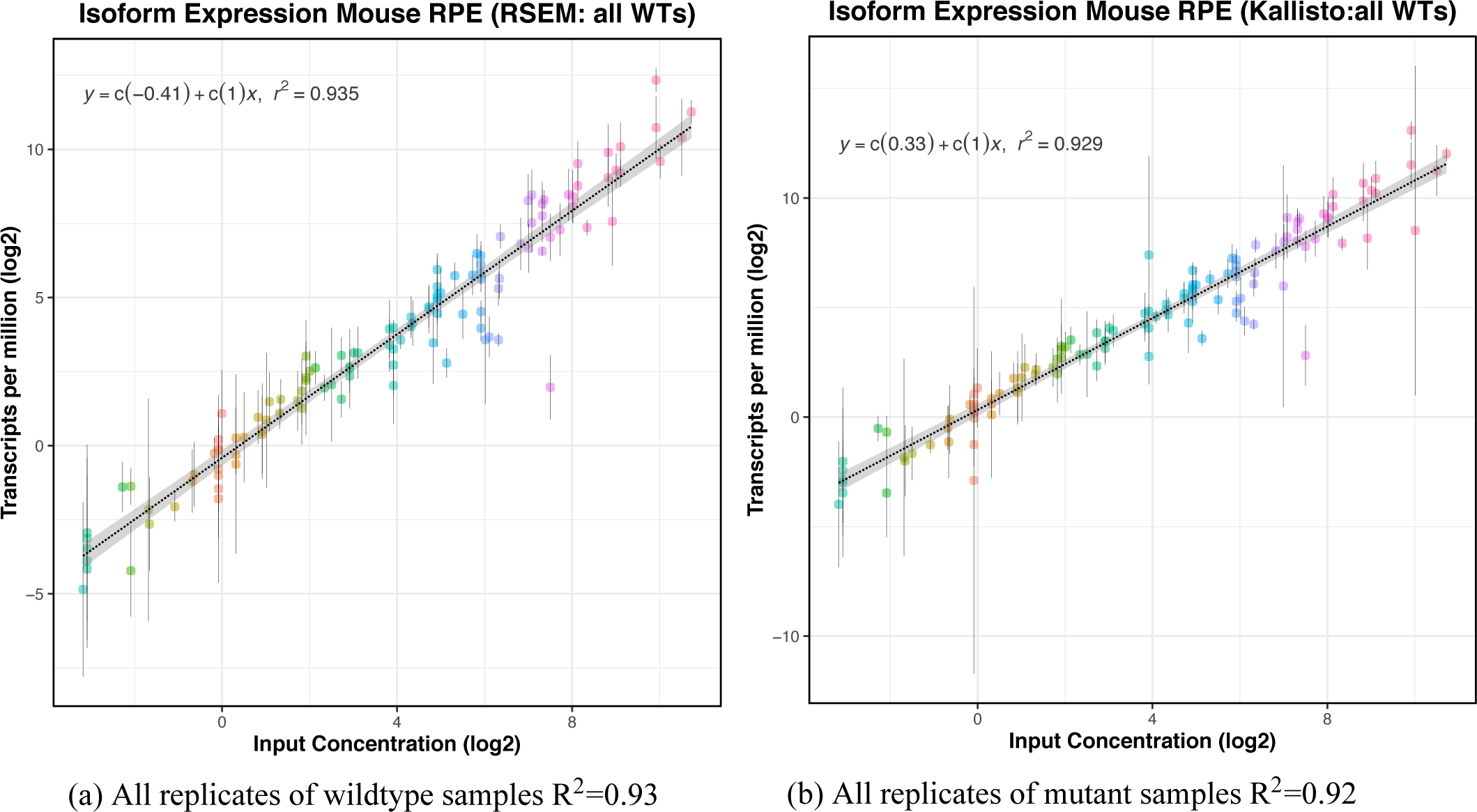
Slope, correlation, LOD and LOQ determination using RSEM and STAR for all the wild type (a) mutant samples (b). Known concentrations are on the x-axis and observed concentration in TPMs is shown on the y-axis. Both the axes are log2 scaled. The vertical bar shows for each sequin isoforms shows the spread of the concentration values. No LOD or LOQ was detected for this sample set.

**Figure (9).**
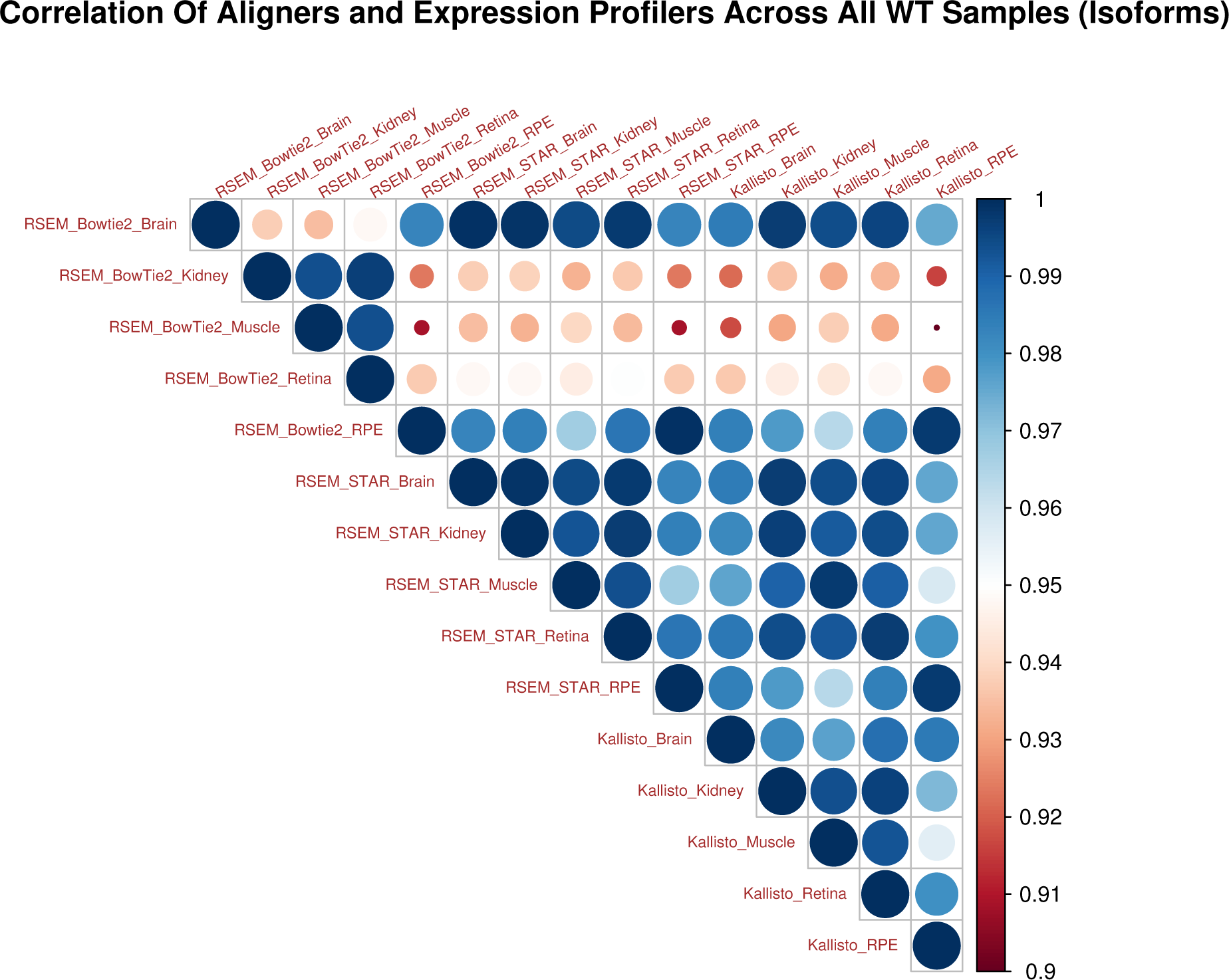
Correlation plot for comparative analysis for all the wild type samples using RSEM and two aligners along with Kallisto across all the mouse tissues. An upper triangular heatmap is generated showing Pearson correlation of determination values using average TPM expression values across the five mouse tissues. The combinations of tools used are shown in the format of:ExpressionTool_Aligner_Tissue, for example, RSEM_STAR_Brain. High correlation of determination values were observed across different tissues types and combinations of different tools. Similar observations were made for the mutant sample set, Supplementary Figure S10.

### Differential Isoform Proportion and Expression

We investigated Differential Transcript Expression (DTE) proportions and directionality of LFC values across all the mouse tissues. As for DGE above, we expect the proportions of the LFC should be similar and unidirectional, provided that the depth of coverage and sequins ratios are proportional across the case and control samples. Supplementary Figure S12 shows proportions and directionality of sequins across all the mouse five tissues.

As with DGE, in most cases unified directionality and proportions of the isoforms was observed. The directionality and proportions of sequins with ground truth of zero LFC are over or under reported by one unit. Additionally, Supplementary Table S11 provides details on some of the specific sequins that were missed by either of both of the DTE tools. Overall, some inconsistencies in reporting of LFC proportions and directionality of the sequins were observed.

Supplementary Figure S11 shows an upper triangular computed correlation of determination in the form of a heatmap using complete case analysis approach. In this case, the missing values (sequins) were removed. The figure shows (R^2^) values for all possible combinations for EBSeq and Sleuth across all the five mouse tissues. Here we followed the “complete case analysis”

Similar to DGE, we know underlying expected fold changes; up-regulation, down-regulation and no change (no LFC) of the sequin isoforms between the case and control sample sets across all the mouse tissues, we can gauge the sensitivity and specificity of various combinations of the tools using receiver operating characteristics (ROC) analysis. Figure 10 shows the ROC curve for all the mouse tissues with different combinations of all the analysis tools.

**Figure (10).**
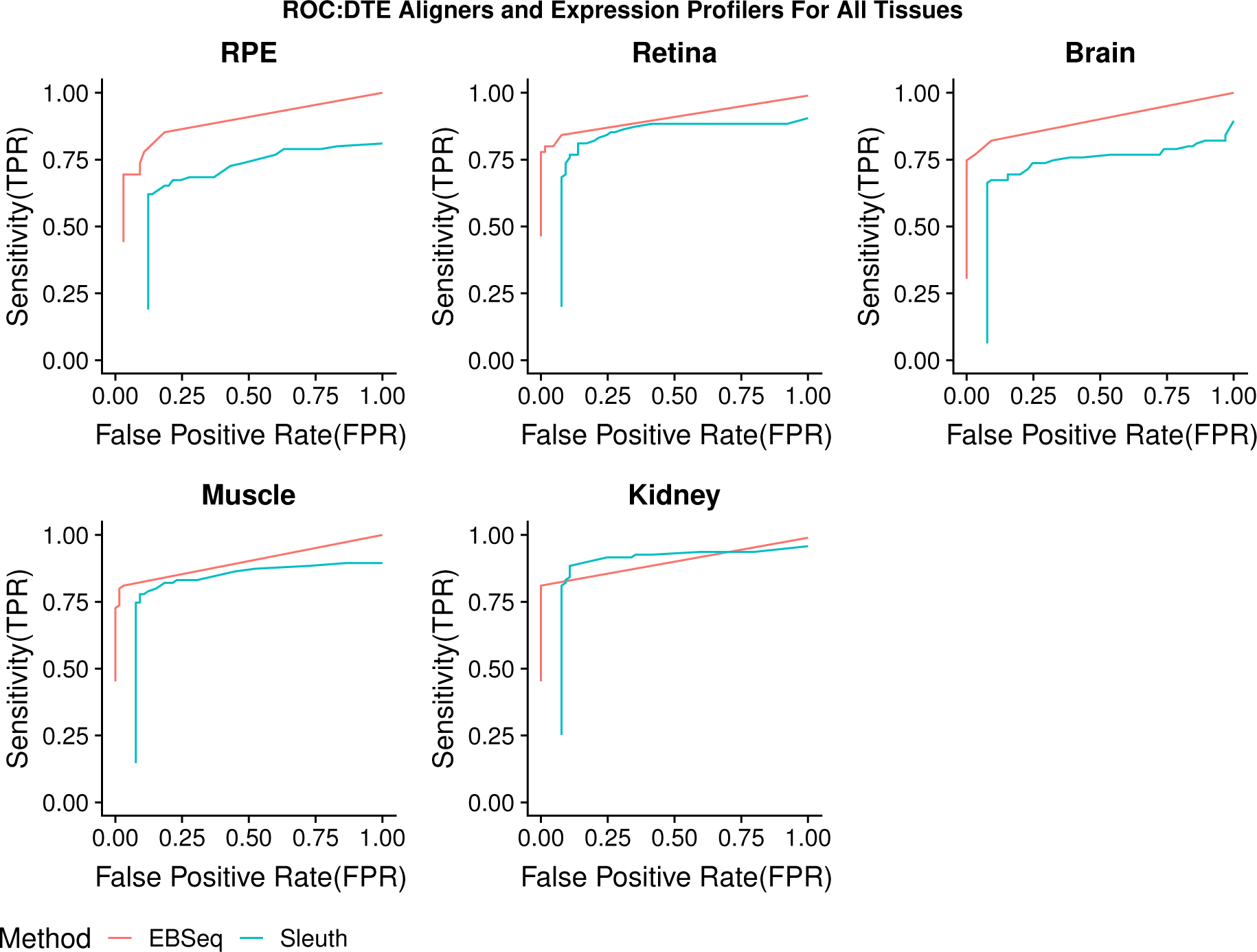
ROC (receiver operating characteristic) curves for Sequin isoforms for diagnostic performance in measuring fold change. Each plot reflects the performance of the aligners, feature read counting and DGE tools across all the five mouse tissues.

Each plot in the figure for a given tissue shows the performance of different combinations of the tools used for the alignments, isoform expression values and DTE. It is evident that the accuracy of isoform detection is not as good as for DGE. The shift in the curves for certain combinations of the tools was due to missed reporting of the sequin isoforms.

### Differential Alternate Splicing

The sequins in each set of samples act as our truth set. Since the amount of each gene and isoform is known *a priori* we can use them for differential alternative splicing (AS) events detection across all the five mouse tissues. Our objective was to compare consistency and reproducibility of each of the splicing detection tools within the samples set and also across the tissue types. The RPE tissue was the first set of samples we used for comparative analysis for differential AS detection tools. Table 2 shows sensitivity for three tools. For RPE, JunctionSeq (86.3%) and MAJIQ (68.18%) outperformed rMATS (31%). Similarly, Table 3 shows specificity with JunctionSeq (97.87%) and MAJIQ (100%) outperforming rMATS(91.48%). We did not use rMATS for further analysis on other tissue types. With the exception of RPE, for the rest of the samples (retina, brain, kidney and muscle), observed sensitivity of JunctionSeq was 100%. MAJIQ showed lower sensitivity ranging from 59% to 90% (Table 2). MAJIQ showed a 100% specificity across all the tissues with JunctionSeq reporting at least one false positive except for the muscle tissue.

**Table (2).** Sequins sensitivity from differential splicing events tools. Calculated sensitivity across all the five mouse tissues is shown. rMATS was tested only on the RPE tissue types. Alignments were made using STAR.

| Tool/<br>Tissue | JunctionSeq |  |  |  |  | MAJIQ |  |  |  |  | rMATS |
| --- | --- | --- | --- | --- | --- | --- | --- | --- | --- | --- | --- |
|  | RPE | Retina | Brain | Muscle | Kidney | RPE | Retina | Brain | Muscle | Kidney | RPE |
| **True Positive** | 19 | 22 | 22 | 22 | 22 | 15 | 19 | 13 | 17 | 20 | 7 |
| **False Negative** | 3 | 0 | 0 | 0 | 0 | 7 | 3 | 9 | 5 | 2 | 15 |
| **Sensitivity%** | 86.3 | 100 | 100 | 100 | 100 | 68.18 | 86.36 | 59.09 | 77.27 | 90.90 | 31.81 |

**Table (3).**
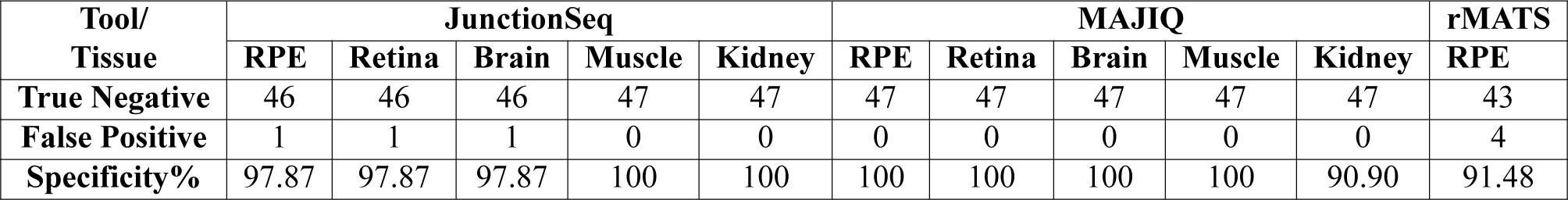
Sequins specificity from differential splicing events tools. Calculated specificity across all the five mouse tissues is shown. rMATS was tested only on the RPE tissue types.

Furthermore, Supplementary Table S12 shows comparative sensitivity from HISAT2 and STAR aligner. In both the case, MAJIQ was used for differential AS detection. HISAT2, in two cases; retina (90%) and brain (63%) slightly outperformed STAR aligner 86% and 59% respectively. In the rest of the samples/tissues, identical results were observed. No true negative was reported.

Additionally, we found comparable results in terms of sensitivity and specificity between the combination of JunctionSeq with STAR and JunctionSeq with HISAT2(Supplementary Table S13) with two exceptions. The combination of HISAT2 and JunctionSeq for retina and muscle samples found to have higher reporting of false positives 5 and 2, respectively than the combination of STAR and JunctionSeq. Near identical sensitivity and specificity was observed the remaining tissue types.

Finally, recall that JunctionSeq is capable of identifying novel splice junctions. We analyzed novel splice junctions reported by JunctionSeq by the two aligners on sequins only. Any reported novel junctions are false positives. Supplementary Table S14 shows comparison of novel splice junction from the two aligners. No more than 3 novel splice junctions per gene was observed. We observed more novel splice junctions from the STAR aligner than HISAT2.

### Impacts of Downsampling on Gene, Isoform and Splicing

We investigated the impacts of lower depth of coverage on detection of gene expression, DGE, isoform expression and DTE and AS events using sequins. We made three tiers of the aligned data set using 150 million, 100 million and 50 million sequence reads. In the previous sections we show that sensitivity and specificity of the aligners, genes and isoform expression profilers are comparable. We used the output from the STAR aligner, featureCount for gene expression profile and used DESeq2 for DGE. Similarly, we used Kallisto for isoform expression, Sleuth for DTE. MAJIQ was used for AS events. Supplementary Table S15, Table S16 and Table S17 show different metrics associated with downsampling for the RPE, retina and brain sample respectively. Various metrics such as, total genomics reads, genomics unique alignment counts, total sequins reads, expected and observed reads counts of sequins post downsampling is shown. Due to lower depth of coverage, two samples in the RPE sample set and one sample in the brain could not be downsampled to 150 million tier. Similarly, two samples in the RPE sample set could not be downsampled to 50 million tier, 5 samples for 100 million and 6 samples for 150 million tier could not downsampled. In all such case the entire sample set was used.

#### Genes

Figure 11 shows the correlation plots of the all three tiers for RPE data set. 2 true positive (TP) sequin was not reported in the 50 million set for RPE. No LOD and LOQ were reported from Anaquin for mouse RPE sample set. Similar observations were made for the retina and brain samples for the all the three tiers. 1 TP sequin was not reported in the 50 milllion set for retina and 1 TP sequins in each of 150 million and 50 million was not reported in the brain tissue. High correlation was observed across between the expected and observed LFC for RPE, retina and brain.

**Figure (11).**
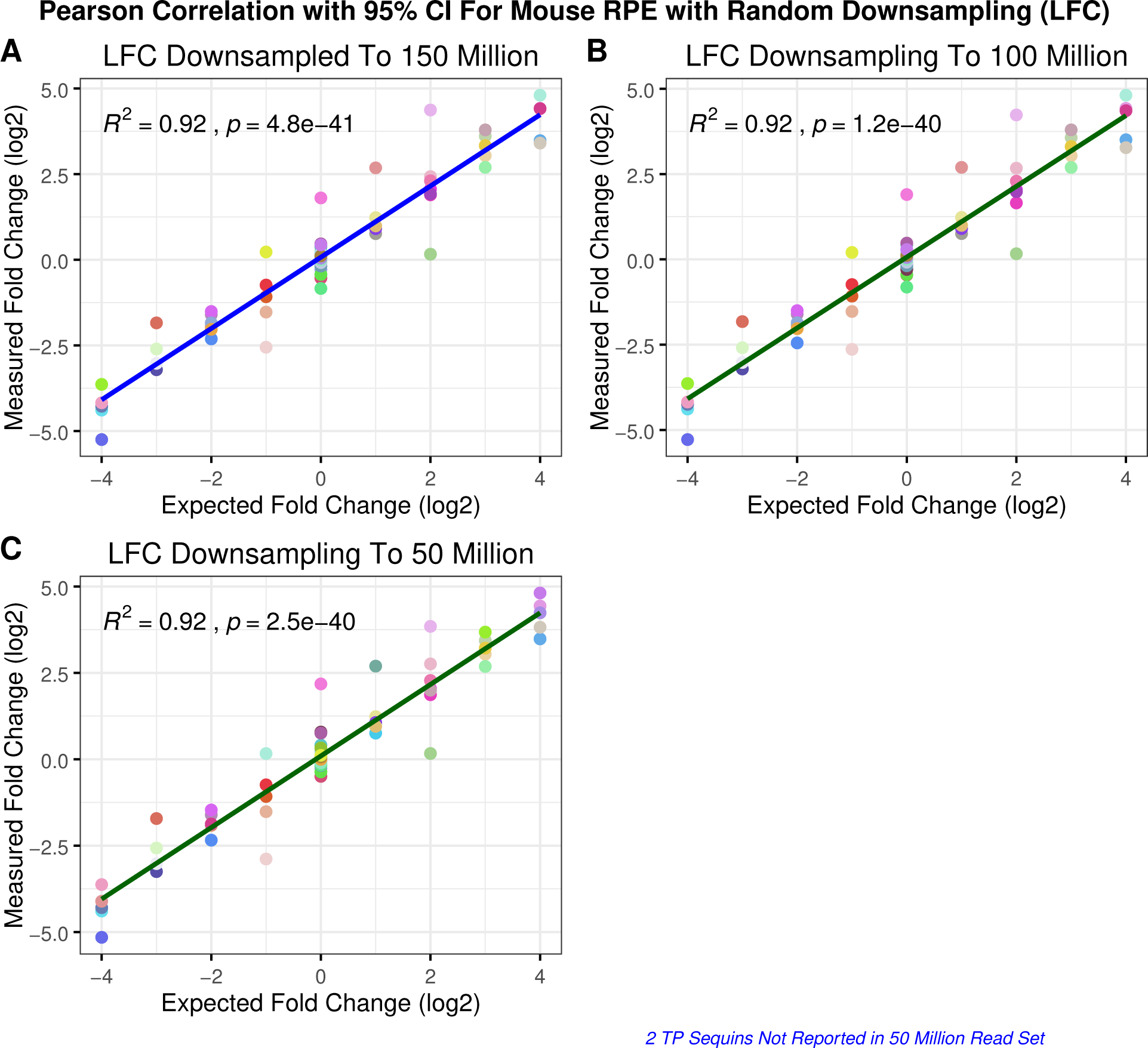
Correlation plot for comparative analysis for mouse RPE for downsampled data. The x-axis represents the expected LFC in log2 values and y-axis represents the observed LFC values by the differential gene expression tools. The plot of LFC for sequin genes across all the three titers is shown. Here we have used DESeq2 only. In the figure, each sequin gene is assigned a unique and static color across the different tissues. The Pearson correlation was calculated for each combinatorial methodology. As per the CI interval, there were some over and some under represented genes. Overall high correlation was observed. We did observed loss of 2 true positive sequins in the 50 million read set.

Next, as with original dataset, we focused on the correlation between the expected and observed LFC values, including the proportions and directionality. Supplementary Figure S13 shows comparison of proportions and directionality for all the three tissues and the three downsampling tiers. As with the complete dataset, some sequins (ranging of 1–2) were not reported. Sequins genes between −1 –+1 showed more variations than the other LFC values.

#### Isoforms

Next we explored impacts of lower DOC on the sequin isoforms. Figure 12, shows correlation between the expected and observed expressions for mouse RPE, tissue with lowest DOC. Similar observations were made with the mutant replicates. No LOD or LOQ was detected for any of the three downsampled tiers and for all three tissues

**Figure (12).**
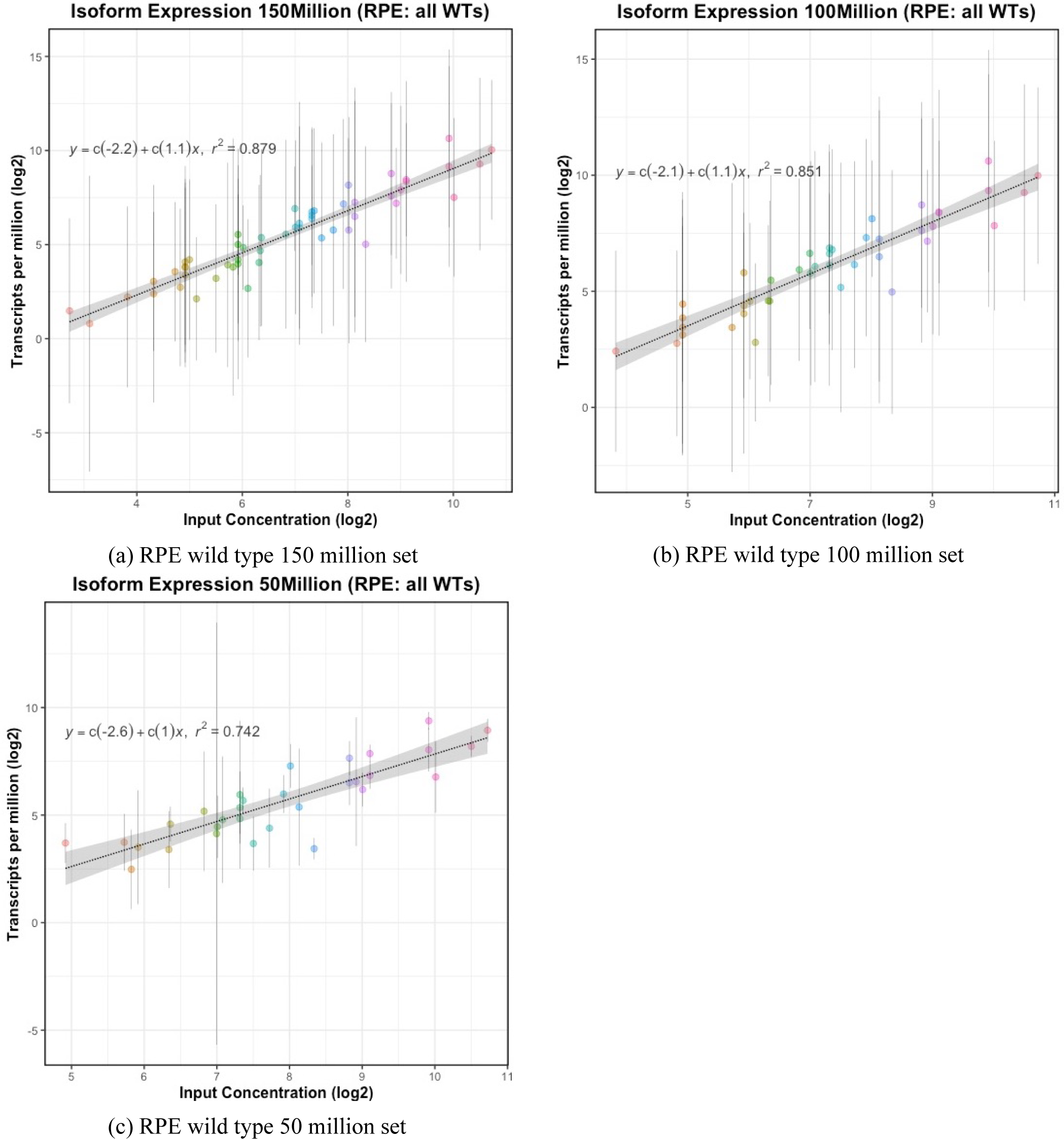
Correlation plot for downsampled mouse RPE wild type (a) mutant tissues (b) for random 150 million reads. Slope, correlation, LOD and LOQ determination for Kallisto is shown. Known concentration are on the x-axis and observed concentration in TPMs is shown on the y-axis. Both the axis are log2 scaled. The vertical bar shows for each sequin isoforms shows the spread of the concentration values. No LOD or LOQ was detected for this sample set. Observed R^2^=0.87 values for WT, R^2^=0.91 for MUT

Table 4 summarizes the observed correlation of determination across brain and retina across the three tiers.

**Table (4).**
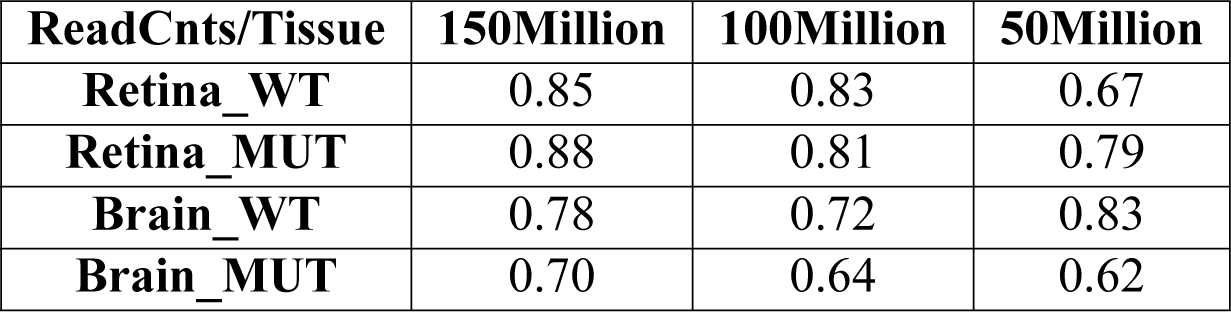
Pearson correlation for downsampled isoforms from mouse retina and brain samples. Random down sampling was performed using different seeds.

Downsampling impacted the isoform LFC the most. No significant LFC sequins were reported for the brain and RPE tissues for all the three titration blocks. In the case of retina, for 100 million reads 6 TP sequins and 1 for the 50 million read set were reported. Similar to isoforms, reporting of AS events also was impacted.

The gene expression and DGE data set remains accurate when the RNA-Seq data are down-sampled. In contrast downsampling noticeability impacted isoform detection and DTE. We observed hight variability across replicates for all the three tiers of the dataset negatively impacting the correlation coefficient.

#### Splicing

Table 5 summarizes the sensitivity of the AS events from MAJIQ. No true negative was reported. For comparison, sensitivity from the original data is also listed. We observed decrease in sensitivity of differential AS detection with lower DOC.

**Table (5).** Sensitivity for detection of AS events on downsampled data. The three mouse tissues were downsampled into 3 tiers; 150, 100 and 50 million read set. MAJIQ was used for AS detection. The reads were aligned using STAR. No true negative was reported. The brain sample set is worst affected. Not that in true sense this is the sample set where each sample set underwent downsampling. In RPE and retina, there were many sample that were used entirely due to lack of read counts.

| **Tissue** | **RPE** |  |  | **Retina** |  |  | **Brain** |  |  |
| --- | --- | --- | --- | --- | --- | --- | --- | --- | --- |
| **Read Cnts** | **150Mill** | **100Mill** | **50Mill** | **150Mill** | **100Mill** | **50Mill** | **150Mill** | **100Mill** | **50Mill** |
| **TP** | 15 | 14 | 13 | 19 | 16 | 15 | 12 | 11 | 9 |
| **FN** | 7 | 8 | 9 | 3 | 6 | 7 | 10 | 11 | 13 |
| **Sensitivity%** | 68.2 | 63.6 | 59.1 | 86.4 | 72.7 | 68.2 | 54.5 | 50.0 | 40.9 |
| **Original Sensitivity%** | 68.18 |  |  | 86.3 |  |  | 59.09 |  |  |

## Discussion

In this study we have evaluated the performance of different tools required to analyze the data from a typical RNA-Seq experiment. We added known mixtures of internal control “sequins” to RNA samples prior to library preparation and sequencing, allowing for quantification of the accuracy of the output from different software tools. Using this approach we demonstrate the use of these internal controls in a RNA-Seq work flow is vital for robust routine assessment of RNA-Seq experiments during experimental and computational development.

Two orthogonal mapping methods, HISAT2 and STAR were evaluated at various levels. We compared unique genomic alignment rate, alignment to synthetic chromosome and to known synthetic intron. In our case, comparable results from both the aligners was observed. We did not observed any preferential bias of one mapper over the other. The performance of the aligner are known to vary based on the genome complexity (different strain and species) [13] and [14]. Our focus remained on sequins validated human and mouse genomes. We did not focus on the run times and there are others studies that have explored this in more details [14] and [45]. Testing every parameter is beyond the scope of this paper. Please see [16] and [13] for more details.

The read count matrices were produced by two different tools, featureCount and HTSeq-Count. This allowed us to perform an unbiased comparison of the features (exons) expression profiles (count matrix) using different combination of the aligners. We did not observed any change in behavior from the aligners or from the expression profile tools. Furthermore, no LOD/LOQ was reported regardless of the aligners, expression count matrix tool and tissue types across the two genotypes and replicates. Expected concentrations of all the genes, inclusive of lowly expressed genes was obtained. Comparable results were obtained from both the expression profilers [25], [23] and [45]. To observe the impacts of lower depth of coverage, we downsampled the original data into three (3) tiers of random read sets. Even with downsampled data, as low as 50 million reads consistent and robust correlation of determination was observed. In all the cases, sensitivity was impacted for genes with should have zero expression. We observed reporting of sequin genes expression in range of −1–+1 units. Such cases reflects both on the alignment, as the reads were aligned to such sequin in a disproportionate manner and further compounded by the expression profilers.

We analyzed DESeq2 and edgeR using different combination of the aligners and read count expression matrix tools. Since sequins were added to all the tissues, this analysis was not only performed in a typical form, that is restricted to within tissues and its corresponding genotypes. The analysis was extended across all the five tissues for comparative analysis. Systematic observations were made: high correlation of determination, consistent proportion and directionality across all the tissues and high sensitivity. We observed that DESeq2 does not report LFC values with low sensitivity, the same could not be said for edgeR. In internal testing, using a third highest LFC value cutoff for edgeR, both the DGE tools were indistinguishable. Both the tools were resilient to any negative impacts of lower depth of coverage. We observed high correlation of determination among the different tissues. For sequin genes which should be constant across the two genotype (sequins mixtures), LFC in range of −1–+1 was reported. In such cases the DGE tools were not able to normalize for disproportionate read alignment and expressions for such specific cases. Using sequins, one can determine the range of LFC that should not be considered and apply such filter to the genomic set to exclude such DGE genes from all downstream analysis.

We analyzed isoform expressions and differential expression using different combinations of the software packages. We used RSEM with Bowtie2 and STAR and another orthogonal software Kallisto for isoform expression within the two genotypes across the five tissues. High correlation was observed with different combination of RSEM and the two aligners. No LOD or LOQ was reported. Similar observation was made with Kallisto. However, robustness of these tools was negatively impacted with lower DOC. We observed a decreasing trend in correlation of determination with corresponding lowering of DOC. Although, with lower DOC, no LOD or LOQ was reported, but extreme variability across the replicates for the sequin isoforms was observed, most with 50 million downsampled dataset. For isoform detection high DOC (>150 million HQ reads) is needed.

EBSeq and Slueth were used for LFC. Combination for RSEM and EBSeq and Kallisto with Sleuth was used. We found irregularities in the reporting of the sequins and their predicted proportions and directionality by both the tools. In almost all the tissues high up-regulated and down-regulated isoforms were reported consistently. The lower expressing sequin isoforms in the range of −1–+1 including isoforms with no (zero) LFC were missed or if reported, inconsistencies in the proportions were observed. DTE tools were intolerant to lower DOC. Insignificant number of true positive sequin isoforms with confidence were reported by the DTE tools. As with isoform expression, caution is advised for DTE with DOC of 150 million reads. The DTE tools were not found as robust and refined as the DGE tools.

We used count-based methods that include both exon-based and event-based approaches; rMATS, MAJIQ and JunctionSeq. We tested alignment from both STAR and HISAT2. The mouse RPE tissue had the lowest depth of coverage compared to other tissues. We first used this tissue to identify best AS tools. MAJIQ and JunctionSeq outperformed rMATS. We next tested and compared MAJIQ and JunctionSeq with rest of the tissues for consistency and reproducibility. Minimal (1) false positives were reported by MAJIQ or JunctionSeq. In most cases with regards to sensitivity and specificity JunctionSeq was found outperforming MAJIQ. In three cases, RPE, retina and brain, JunctionSeq reported one false positive, none was reported by MAJIQ.

Since absence of reporting of AS events in classical sense, we found it hard to interpret the observed phenomenas from JunctionSeq used as a proxy to AS or convert the output to be more interpretable compared to MAJIQ. Even though JunctionSeq produces a number of images for DEU, we found that output from MAJIQ’s companion tool “Volia” was easier to interpret. With such a high depth of coverage we find it disconcerting that not all the true positive sequins were reported by MAJIQ. Lower depth of coverage impacted sensitivity to predict AS events of known true positive events. A decreasing trend is the sensitivity across three tiers of downsampling and three tissue was observed. No true negative event was reported.

We used sequins as ground truth for systematic evaluate different tools used during RNA-Seq analysis. We demonstrated that inclusion of internal controls in RNA-seq experiments allows accurate determination of detection levels, and better assessment of DGE, DTE and AS accuracy. The gene expression profiling and DGE tools are found to be robust. The isoform expression profiling tools were robust, but not as tolerant to lower depth of coverage as the DGE tools. We recommend high depth of coverage, about 200 million reads for confident prediction of differential isoform expression. Even with high DOC, for AS event detection, we found the current tools lacking high sensitivity. This was compounded by lower depth of coverage. More efforts are needed to improve specificity and sensitivity of DTE and AS detection.

## Supplementary Material

**Table (S1).** Genomes and versions. Table lists all the genomes and corresponding versions used in the analysis.

| Reference | Version |
| --- | --- |
| Mouse Genome | MM10 |
| Mouse GENCODE Gene Model | M12 (Ensembl 87) |
| Mouse rRNA NCBI | Nucleotide: txid10090 |

**Table (S2).** Analysis tools and versions. Table lists all the analysis tools that were used during the analysis.

| Software | Version |
| --- | --- |
| FastQC | 0.11.3 |
| MultiQC | 1.2 |
| STAR | 2.5.3a |
| HISAT2 | 2.1.0 |
| rMATS (turbo) | 4.0.2 |
| MAJIQ | 1.1.5 |
| SubRead | 1.5.2 |
| HTSeq | 0.9.1 |
| RSeQC | 2.6.4 |
| edgeR | 3.22.2 |
| DESeq2 | 1.20.0 |
| SmartSVA | 0.1.3 |
| SVA | 0.1.3 |
| Samtools | 1.4 |
| BowTie2 | 2.2.4 |
| Picardtools | 1.87 |
| Anaquin | 2.4.0 |
| RStudio | 1.1.453 |
| R | 3.5.0 |
| Corrplot | 0.85 |

**Figure (S1).**
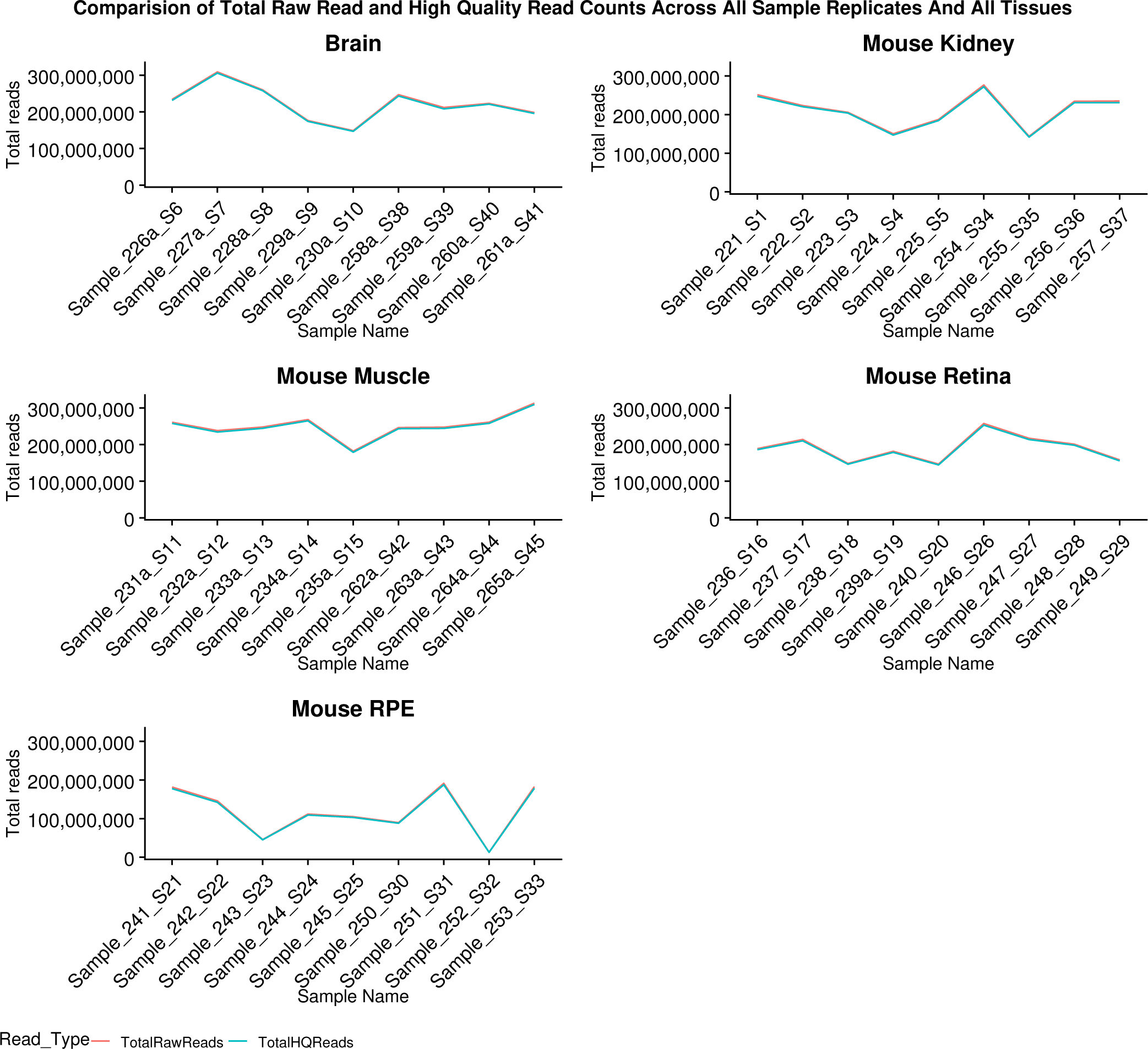
Total read comparative pre and post processing. The figure shows total raw reads for each sample replicates including reads counts post quality processing. Total reads retained post QC ranged from 98%-99%, reflective in indistinguishable trend lines. The muscle and brain sample sets were among the highest read counts and as expected the RPE sample set showed the lowest read counts. For visualization only, discrete data is shown in a continuous form. From left to right, first 5 are mutants followed by wild type samples. HQ refers to reads and bases that have passed strict QC/QA processes.

**Table (S3).** Summary statistics of the first sequencing run. Table lists various standard summary statistics. RC refers to read counts and SRun refers to sequencing run.

| Tissue | Min_RC | Max_RC | Mean | q1 | q3 | sstdev | SRun |
| --- | --- | --- | --- | --- | --- | --- | --- |
| Brain_MUT | 17,774,374.00 | 47,831,123.00 | 32,978,284.80 | 30,247,107.00 | 38,115,267.00 | 11,066,114.35 | 180316_SideA |
| Brain_WT | 44,964,350.00 | 53,334,326.00 | 49,872,655.25 | 48,962,570.75 | 51,506,057.00 | 3,526,178.06 | 180316_SideA |
| Kidney_MUT | 18,908,291.00 | 61,991,216.00 | 40,235,123.20 | 33,223,708.00 | 47,277,456.00 | 16,024,415.00 | 180316_SideA |
| Kidney_WT | 29,640,312.00 | 50,646,709.00 | 38,913,504.75 | 34,499,694.00 | 42,097,309.75 | 8,785,940.94 | 180316_SideA |
| Muscle_MUT | 33,060,197.00 | 64,369,275.00 | 49,152,008.00 | 45,138,649.00 | 56,726,582.00 | 11,950,010.66 | 180316_SideA |
| Muscle_WT | 38,495,751.00 | 79,720,781.00 | 58,744,616.50 | 47,683,314.00 | 69,442,269.50 | 17,952,446.39 | 180316_SideA |
| RPE_MUT | 10,155,891.00 | 26,240,789.00 | 20,287,271.80 | 18,934,136.00 | 25,800,007.00 | 6,525,159.07 | 180316_SideA |
| RPE_WT | 2,322,331.00 | 53,878,507.00 | 29,086,839.00 | 13,988,974.75 | 45,171,123.25 | 23,312,138.81 | 180316_SideA |
| Retina_MUT | 29,616,145.00 | 53,113,782.00 | 39,190,498.00 | 32,174,368.00 | 45,054,179.00 | 9,738,485.57 | 180316_SideA |
| Retina_WT | 22,053,007.00 | 45,558,421.00 | 34,166,835.00 | 29,885,213.50 | 38,809,577.50 | 9,747,341.21 | 180316_SideA |

**Table (S4).** Summary statistics of the second sequencing run. Table lists various standard summary statistics. RC refers to read counts and SRun refers to sequencing run.

| Tissue | Min_RC | Max_RC | Mean | q1 | q3 | sstdev | SRun |
| --- | --- | --- | --- | --- | --- | --- | --- |
| Brain_MUT | 32,152,639.00 | 72,220,552.00 | 48,857,344.80 | 38,315,775.00 | 54,289,238.00 | 15,555,671.49 | 180329_SideB |
| Brain_WT | 29,380,289.00 | 50,532,898.00 | 41,078,208.50 | 33,540,529.25 | 49,737,502.75 | 10,559,804.43 | 180329_SideB |
| Kidney_MUT | 29,168,341.00 | 48,734,586.00 | 41,962,012.20 | 39,821,176.00 | 47,259,649.00 | 7,911,986.32 | 180329_SideB |
| Kidney_WT | 30,573,417.00 | 56,677,704.00 | 49,143,940.50 | 48,246,328.50 | 55,559,932.50 | 12,424,146.74 | 180329_SideB |
| Muscle_MUT | 39,911,361.00 | 55,456,660.00 | 48,490,047.20 | 46,087,495.00 | 52,144,800.00 | 5,945,303.49 | 180329_SideB |
| Muscle_WT | 48,308,605.00 | 72,138,103.00 | 55,958,736.00 | 49,415,781.25 | 58,237,072.75 | 11,014,551.57 | 180329_SideB |
| RPE_MUT | 8,690,207.00 | 40,478,959.00 | 25,355,064.40 | 20,918,726.00 | 34,186,494.00 | 12,372,054.76 | 180329_SideB |
| RPE_WT | 2,691,785.00 | 37,939,584.00 | 23,939,273.00 | 14,265,830.00 | 37,236,304.50 | 16,851,461.92 | 180329_SideB |
| Retina_MUT | 19,429,820.00 | 45,695,610.00 | 31,733,182.80 | 25,549,296.00 | 34,967,584.00 | 9,958,527.80 | 180329_SideB |
| Retina_WT | 30,775,728.00 | 53,812,548.00 | 45,462,640.50 | 43,368,478.50 | 50,725,305.00 | 10,128,737.42 | 180329_SideB |

**Table (S5).**
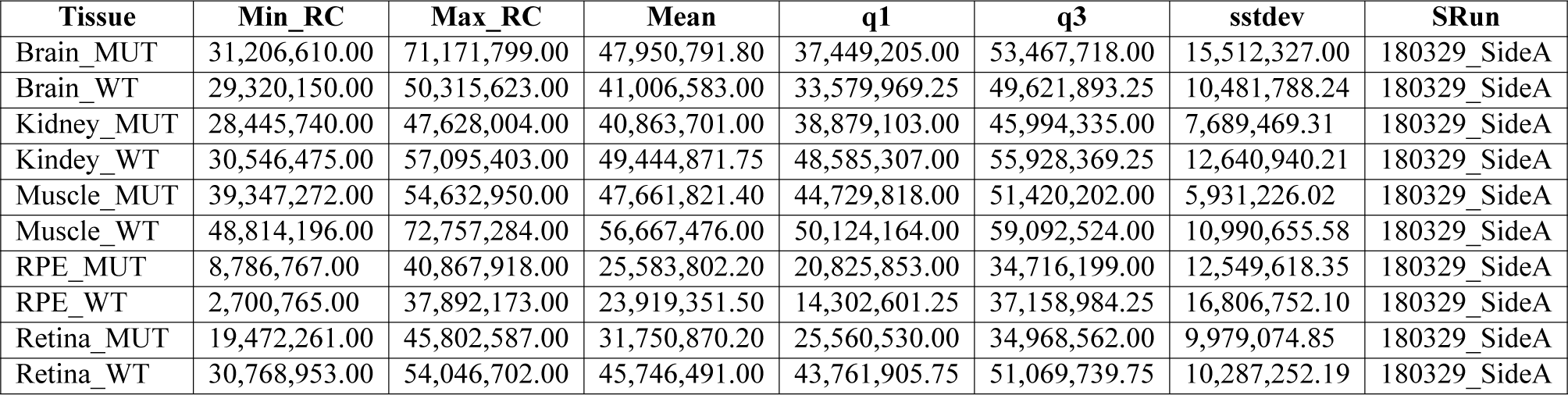
Summary statistics of the third sequencing run. Table lists various standard summary statistics. RC refers to read counts and SRun refers to sequencing run.

| Tissue | Min_RC | Max_RC | Mean | q1 | q3 | sstdev | SRun |
| --- | --- | --- | --- | --- | --- | --- | --- |
| Brain_MUT | 31,206,610.00 | 71,171,799.00 | 47,950,791.80 | 37,449,205.00 | 53,467,718.00 | 15,512,327.00 | 180329_SideA |
| Brain_WT | 29,320,150.00 | 50,315,623.00 | 41,006,583.00 | 33,579,969.25 | 49,621,893.25 | 10,481,788.24 | 180329_SideA |
| Kidney_MUT | 28,445,740.00 | 47,628,004.00 | 40,863,701.00 | 38,879,103.00 | 45,994,335.00 | 7,689,469.31 | 180329_SideA |
| Kidney_WT | 30,546,475.00 | 57,095,403.00 | 49,444,871.75 | 48,585,307.00 | 55,928,369.25 | 12,640,940.21 | 180329_SideA |
| Muscle_MUT | 39,347,272.00 | 54,632,950.00 | 47,661,821.40 | 44,729,818.00 | 51,420,202.00 | 5,931,226.02 | 180329_SideA |
| Muscle_WT | 48,814,196.00 | 72,757,284.00 | 56,667,476.00 | 50,124,164.00 | 59,092,524.00 | 10,990,655.58 | 180329_SideA |
| RPE_MUT | 8,786,767.00 | 40,867,918.00 | 25,583,802.20 | 20,825,853.00 | 34,716,199.00 | 12,549,618.35 | 180329_SideA |
| RPE_WT | 2,700,765.00 | 37,892,173.00 | 23,919,351.50 | 14,302,601.25 | 37,158,984.25 | 16,806,752.10 | 180329_SideA |
| Retina_MUT | 19,472,261.00 | 45,802,587.00 | 31,750,870.20 | 25,560,530.00 | 34,968,562.00 | 9,979,074.85 | 180329_SideA |
| Retina_WT | 30,768,953.00 | 54,046,702.00 | 45,746,491.00 | 43,761,905.75 | 51,069,739.75 | 10,287,252.19 | 180329_SideA |

**Table (S6).** Summary statistics of the fourth sequencing run. Table lists various standard summary statistics. RC refers to read counts and SRun refers to sequencing run.

| Tissue | Min_RC | Max_RC | Mean | q1 | q3 | sstdev | SRun |
| --- | --- | --- | --- | --- | --- | --- | --- |
| Brain_MUT | 27,575,093.00 | 63,976,417.00 | 48,166,448.40 | 35,337,636.00 | 61,440,168.00 | 16,074,312.89 | 180420_SideB |
| Brain_WT | 36,470,873.00 | 50,193,507.00 | 43,581,499.50 | 41,881,761.50 | 45,530,547.00 | 5,610,889.88 | 180420_SideB |
| Kidney_MUT | 33,457,303.00 | 48,259,293.00 | 40,589,584.20 | 37,066,118.00 | 43,924,826.00 | 5,773,960.14 | 180420_SideB |
| Kidney_WT | 23,225,180.00 | 66,183,582.00 | 42,328,409.75 | 33,320,651.00 | 48,960,197.25 | 17,950,254.02 | 180420_SideB |
| Muscle_MUT | 29,438,918.00 | 57,359,123.00 | 47,734,526.00 | 48,031,299.00 | 52,781,014.00 | 10,770,227.19 | 180420_SideB |
| Muscle_WT | 41,873,226.00 | 50,643,895.00 | 47,418,327.75 | 45,894,790.50 | 50,101,632.25 | 3,976,962.41 | 180420_SideB |
| RPE_MUT | 9,074,588.00 | 37,295,963.00 | 23,545,374.60 | 22,435,199.00 | 25,669,359.00 | 10,053,759.91 | 180420_SideB |
| RPE_WT | 2,631,976.00 | 36,212,909.00 | 20,968,421.75 | 14,082,932.50 | 29,399,890.25 | 14,329,301.50 | 180420_SideB |
| Retina_MUT | 29,473,722.00 | 45,367,879.00 | 36,814,789.00 | 30,069,562.00 | 39,958,973.00 | 6,858,170.49 | 180420_SideB |
| Retina_WT | 25,721,447.00 | 62,597,155.00 | 41,684,266.50 | 32,352,273.50 | 48,541,225.00 | 15,786,021.55 | 180420_SideB |

**Table (S7).** Summary statistics of the fifth sequencing run. Table lists various standard summary statistics. RC refers to read counts and SRun refers to sequencing run.

| Tissue | Min_RC | Max_RC | Mean | q1 | q3 | sstdev | SRun |
| --- | --- | --- | --- | --- | --- | --- | --- |
| Brain_MUT | 26,876,041.00 | 63,668,498.00 | 47,740,737.40 | 34,978,434.00 | 61,202,734.00 | 16,210,221.15 | 180420_SideA |
| Brain_WT | 36,918,458.00 | 51,027,046.00 | 44,568,074.25 | 43,034,603.00 | 46,696,867.75 | 5,801,149.03 | 180420_SideA |
| Kidney_MUT | 32,463,591.00 | 47,972,862.00 | 40,001,739.60 | 36,662,935.00 | 43,323,351.00 | 5,973,191.54 | 180420_SideA |
| Kidney_WT | 23,221,786.00 | 66,953,762.00 | 42,621,368.75 | 33,643,813.75 | 49,132,518.50 | 18,248,506.12 | 180420_SideA |
| Muscle_MUT | 28,622,627.00 | 55,655,034.00 | 46,419,083.40 | 46,707,669.00 | 51,558,077.00 | 10,468,099.28 | 180420_SideA |
| Muscle_WT | 42,425,732.00 | 51,545,746.00 | 48,249,609.75 | 46,647,989.75 | 51,115,100.50 | 4,172,479.27 | 180420_SideA |
| RPE_MUT | 9,044,078.00 | 36,818,958.00 | 23,236,333.40 | 21,887,215.00 | 25,449,486.00 | 9,908,212.38 | 180420_SideA |
| RPE_WT | 2,672,613.00 | 37,131,206.00 | 21,277,501.50 | 13,818,348.75 | 30,112,246.25 | 14,761,265.52 | 180420_SideA |
| Retina_MUT | 29,204,397.00 | 44,957,206.00 | 36,465,152.40 | 29,553,853.00 | 39,923,484.00 | 6,882,867.79 | 180420_SideA |
| Retina_WT | 25,431,971.00 | 62,675,224.00 | 41,452,999.50 | 32,082,930.50 | 48,222,470.50 | 15,937,718.54 | 180420_SideA |

**Figure (S2).**
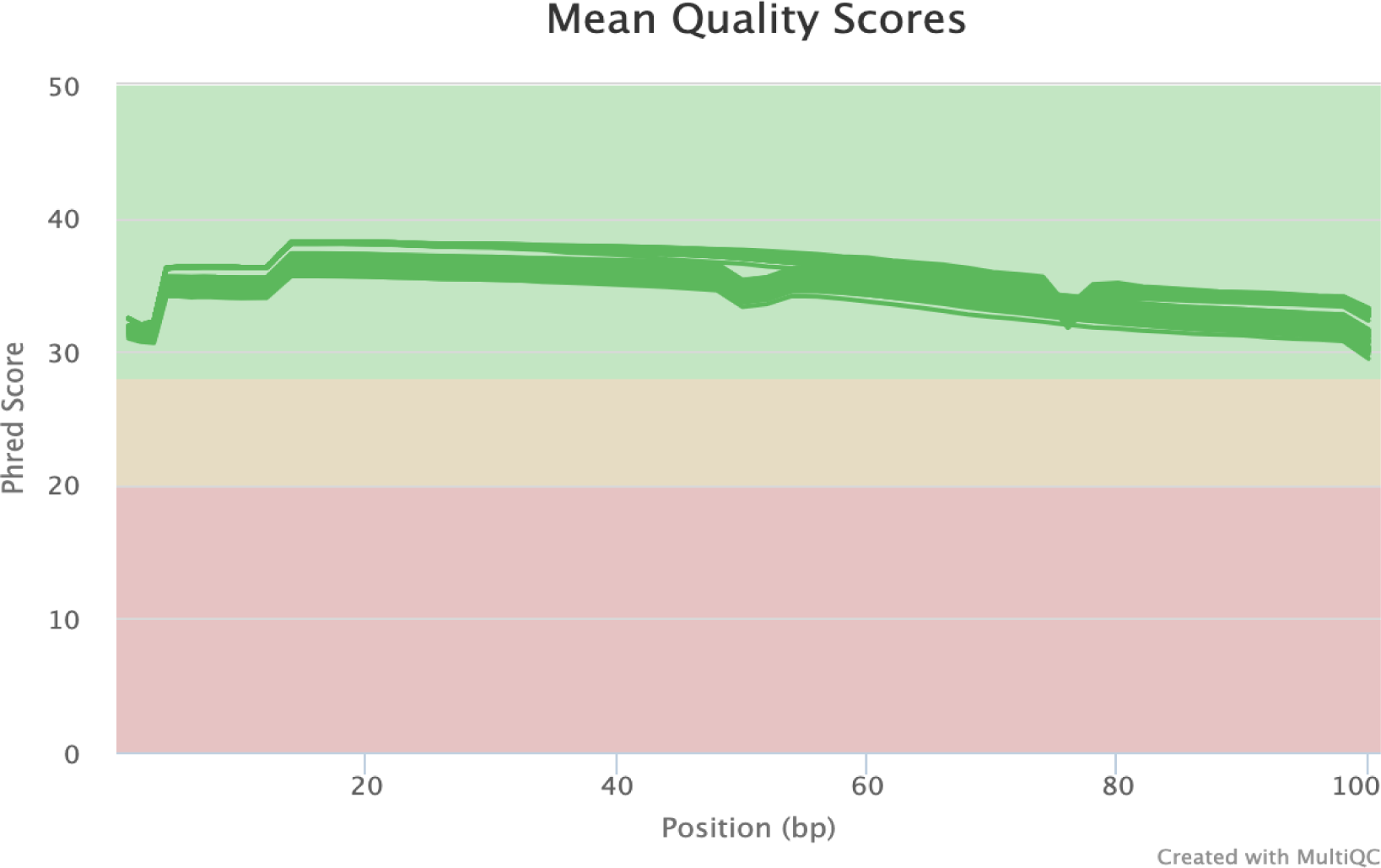
Comparison of total reads of all the 45 samples. High mean phred quality per base, per cycle for the entire read length was observed. No biases were observed.

**Table (S8).** Sequins for differential genes expression. In the table all the true positive genes from the two mixtures are shown in attomoles/*µ*l units, sorted from minimum-maximum. The genes cover true positive (TP), upregulation, downregulation and uniform (no change) set of sequins in the data set. There are 76 sequins genes for DGE.

| **ID** | **MixA** | **MixB** | **ID** | **MixA** | **MixB** |
| --- | --- | --- | --- | --- | --- |
| R1_51 | 0.059 | 0.059 | R2_63 | 1.8883 | 1.8883 |
| R2_76 | 0.059 | 0.059 | R2_73 | 1.8883 | 3.7766 |
| R2_33 | 0.059 | 0.118 | R1_11 | 1933.59 | 1933.59 |
| R1_24 | 0.059 | 0.236 | R2_154 | 1933.59 | 1933.59 |
| R1_21 | 0.059 | 0.4721 | R2_72 | 1933.59 | 1933.59 |
| R2_105 | 0.118 | 0.059 | R1_31 | 241.699 | 241.699 |
| R1_93 | 0.118 | 0.118 | R2_46 | 241.699 | 241.699 |
| R2_152 | 0.118 | 0.118 | R1_73 | 241.699 | 483.398 |
| R2_47 | 0.118 | 0.118 | R2_41 | 241.699 | 60.4248 |
| R2_115 | 0.118 | 0.236 | R1_91 | 30.2124 | 120.85 |
| R1_53 | 0.236 | 0.059 | R2_32 | 30.2124 | 15.1062 |
| R2_37 | 0.236 | 0.059 | R1_23 | 30.2124 | 30.2124 |
| R1_63 | 0.236 | 0.236 | R1_71 | 30.2124 | 30.2124 |
| R2_54 | 0.236 | 0.236 | R2_116 | 3.73883 | 3.73883 |
| R1_82 | 0.236 | 1.8883 | R2_68 | 3.7766 | 0.236 |
| R1_42 | 0.4721 | 0.4721 | R1_62 | 3.7766 | 0.9441 |
| R1_81 | 0.4721 | 0.4721 | R2_67 | 3.7766 | 15.1062 |
| R2_20 | 0.4721 | 1.8883 | R2_71 | 3.7766 | 3.7766 |
| R1_12 | 0.4721 | 7.5531 | R2_6 | 483.398 | 241.699 |
| R2_143 | 0.4721 | 7.5531 | R1_22 | 483.398 | 30.2124 |
| R2_153 | 0.9441 | 0.4721 | R2_38 | 483.398 | 483.398 |
| R1_83 | 0.9441 | 0.9441 | R2_42 | 483.398 | 483.398 |
| R2_1 | 0.9441 | 0.9441 | R1_14 | 483.398 | 966.797 |
| R1_52 | 0.9441 | 15.1062 | R2_140 | 60.4248 | 30.2124 |
| R1_43 | 0.9441 | 7.5531 | R1_32 | 60.4248 | 3.7766 |
| R1_33 | 120.85 | 120.85 | R2_117 | 60.4248 | 60.4248 |
| R1_41 | 120.85 | 241.699 | R2_28 | 60.4248 | 60.4248 |
| R1_92 | 120.85 | 30.2124 | R2_26 | 60.4248 | 7.5531 |
| R1_13 | 120.85 | 483.398 | R2_27 | 7.5531 | 0.9441 |
| R2_53 | 120.85 | 966.797 | R2_57 | 7.5531 | 0.9441 |
| R2_24 | 15.1062 | 15.1062 | R2_59 | 7.5531 | 120.85 |
| R2_45 | 15.1062 | 15.1062 | R2_60 | 7.5531 | 120.85 |
| R1_101 | 15.1062 | 1.8883 | R1_61 | 7.5531 | 7.5531 |
| R2_19 | 15.1062 | 3.7766 | R2_150 | 966.797 | 120.85 |
| R1_102 | 15.1062 | 60.4248 | R1_103 | 966.797 | 483.398 |
| R1_72 | 1.8883 | 0.118 | R2_7 | 966.797 | 60.4248 |
| R2_55 | 1.8883 | 0.4721 | R2_14 | 966.797 | 966.797 |
| R2_151 | 1.8883 | 1.8883 | R2_65 | 966.797 | 966.797 |

**Table (S9).**
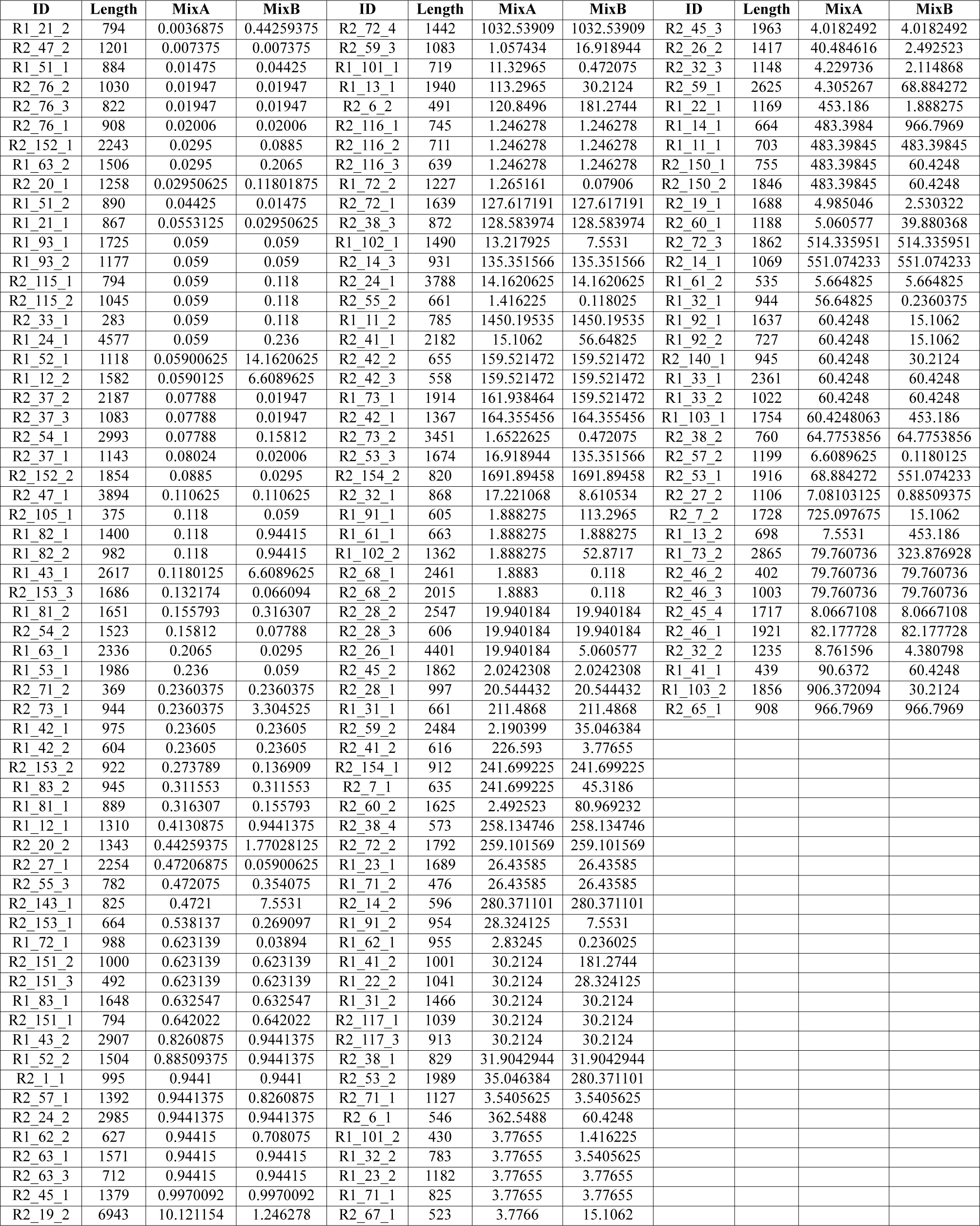
Sequins for differential isoform expression. In the table TP isoforms of the two mixtures (in attomoles/*µ*l units) along with the length are shown, sorted from minimum-maximum of their respective concentration. The isoforms cover upregulation, downregulation and uniform (no change) set of sequins in the data set. There are 160 sequins isoform counts.

| ID | Length | MixA | MixB | ID | Length | MixA | MixB | ID | Length | MixA | MixB |
| --- | --- | --- | --- | --- | --- | --- | --- | --- | --- | --- | --- |
| R1_21_2 | 794 | 0.0036875 | 0.44259375 | R2_72_4 | 1442 | 1032.53909 | 1032.53909 | R2_45_3 | 1963 | 4.0182492 | 4.0182492 |
| R2_47_2 | 1201 | 0.007375 | 0.007375 | R2_59_3 | 1083 | 1.057434 | 16.918944 | R2_26_2 | 1417 | 40.484616 | 2.492523 |
| R1_51_1 | 884 | 0.01475 | 0.04425 | R1_101_1 | 719 | 11.32965 | 0.472075 | R2_32_3 | 1148 | 4.229736 | 2.114868 |
| R2_76_2 | 1030 | 0.01947 | 0.01947 | R1_13_1 | 1940 | 113.2965 | 30.2124 | R2_59_1 | 2625 | 4.305267 | 68.884272 |
| R2_76_3 | 822 | 0.01947 | 0.01947 | R2_6_2 | 491 | 120.8496 | 181.2744 | R1_22_1 | 1169 | 453.186 | 1.888275 |
| R2_76_1 | 908 | 0.02006 | 0.02006 | R2_116_1 | 745 | 1.246278 | 1.246278 | R1_14_1 | 664 | 483.3984 | 966.7969 |
| R2_152_1 | 2243 | 0.0295 | 0.0885 | R2_116_2 | 711 | 1.246278 | 1.246278 | R1_11_1 | 703 | 483.39845 | 483.39845 |
| R1_63_2 | 1506 | 0.0295 | 0.2065 | R2_116_3 | 639 | 1.246278 | 1.246278 | R2_150_1 | 755 | 483.39845 | 60.4248 |
| R2_20_1 | 1258 | 0.02950625 | 0.11801875 | R1_72_2 | 1227 | 1.265161 | 0.07906 | R2_150_2 | 1846 | 483.39845 | 60.4248 |
| R1_51_2 | 890 | 0.04425 | 0.01475 | R2_72_1 | 1639 | 127.617191 | 127.617191 | R2_19_1 | 1688 | 4.985046 | 2.530322 |
| R1_21_1 | 867 | 0.0553125 | 0.02950625 | R2_38_3 | 872 | 128.583974 | 128.583974 | R2_60_1 | 1188 | 5.060577 | 39.880368 |
| R1_93_1 | 1725 | 0.059 | 0.059 | R1_102_1 | 1490 | 13.217925 | 7.5531 | R2_72_3 | 1862 | 514.335951 | 514.335951 |
| R1_93_2 | 1177 | 0.059 | 0.059 | R2_14_3 | 931 | 135.351566 | 135.351566 | R2_14_1 | 1069 | 551.074233 | 551.074233 |
| R2_115_1 | 794 | 0.059 | 0.118 | R2_24_1 | 3788 | 14.1620625 | 14.1620625 | R1_61_2 | 535 | 5.664825 | 5.664825 |
| R2_115_2 | 1045 | 0.059 | 0.118 | R2_55_2 | 661 | 1.416225 | 0.118025 | R1_32_1 | 944 | 56.64825 | 0.2360375 |
| R2_33_1 | 283 | 0.059 | 0.118 | R1_11_2 | 785 | 1450.19535 | 1450.19535 | R1_92_1 | 1637 | 60.4248 | 15.1062 |
| R1_24_1 | 4577 | 0.059 | 0.236 | R2_41_1 | 2182 | 15.1062 | 56.64825 | R1_92_2 | 727 | 60.4248 | 15.1062 |
| R1_52_1 | 1118 | 0.05900625 | 14.1620625 | R2_42_2 | 655 | 159.521472 | 159.521472 | R2_140_1 | 945 | 60.4248 | 30.2124 |
| R1_12_2 | 1582 | 0.0590125 | 6.6089625 | R2_42_3 | 558 | 159.521472 | 159.521472 | R1_33_1 | 2361 | 60.4248 | 60.4248 |
| R2_37_2 | 2187 | 0.07788 | 0.01947 | R1_73_1 | 1914 | 161.938464 | 159.521472 | R1_33_2 | 1022 | 60.4248 | 60.4248 |
| R2_37_3 | 1083 | 0.07788 | 0.01947 | R2_42_1 | 1367 | 164.355456 | 164.355456 | R1_103_1 | 1754 | 60.4248063 | 453.186 |
| R2_54_1 | 2993 | 0.07788 | 0.15812 | R2_73_2 | 3451 | 1.6522625 | 0.472075 | R2_38_2 | 760 | 64.7753856 | 64.7753856 |
| R2_37_1 | 1143 | 0.08024 | 0.02006 | R2_53_3 | 1674 | 16.918944 | 135.351566 | R2_57_2 | 1199 | 6.6089625 | 0.1180125 |
| R2_152_2 | 1854 | 0.0885 | 0.0295 | R2_154_2 | 820 | 1691.89458 | 1691.89458 | R2_53_1 | 1916 | 68.884272 | 551.074233 |
| R2_47_1 | 3894 | 0.110625 | 0.110625 | R2_32_1 | 868 | 17.221068 | 8.610534 | R2_27_2 | 1106 | 7.08103125 | 0.88509375 |
| R2_105_1 | 375 | 0.118 | 0.059 | R1_91_1 | 605 | 1.888275 | 113.2965 | R2_7_2 | 1728 | 725.097675 | 15.1062 |
| R1_82_1 | 1400 | 0.118 | 0.94415 | R1_61_1 | 663 | 1.888275 | 1.888275 | R1_13_2 | 698 | 7.5531 | 453.186 |
| R1_82_2 | 982 | 0.118 | 0.94415 | R1_102_2 | 1362 | 1.888275 | 52.8717 | R1_73_2 | 2865 | 79.760736 | 323.876928 |
| R1_43_1 | 2617 | 0.1180125 | 6.6089625 | R2_68_1 | 2461 | 1.8883 | 0.118 | R2_46_2 | 402 | 79.760736 | 79.760736 |
| R2_153_3 | 1686 | 0.132174 | 0.066094 | R2_68_2 | 2015 | 1.8883 | 0.118 | R2_46_3 | 1003 | 79.760736 | 79.760736 |
| R1_81_2 | 1651 | 0.155793 | 0.316307 | R2_28_2 | 2547 | 19.940184 | 19.940184 | R2_45_4 | 1717 | 8.0667108 | 8.0667108 |
| R2_54_2 | 1523 | 0.15812 | 0.07788 | R2_28_3 | 606 | 19.940184 | 19.940184 | R2_46_1 | 1921 | 82.177728 | 82.177728 |
| R1_63_1 | 2336 | 0.2065 | 0.0295 | R2_26_1 | 4401 | 19.940184 | 5.060577 | R2_32_2 | 1235 | 8.761596 | 4.380798 |
| R1_53_1 | 1986 | 0.236 | 0.059 | R2_45_2 | 1862 | 2.0242308 | 2.0242308 | R1_41_1 | 439 | 90.6372 | 60.4248 |
| R2_71_2 | 369 | 0.2360375 | 0.2360375 | R2_28_1 | 997 | 20.544432 | 20.544432 | R1_103_2 | 1856 | 906.372094 | 30.2124 |
| R2_73_1 | 944 | 0.2360375 | 3.304525 | R1_31_1 | 661 | 211.4868 | 211.4868 | R2_65_1 | 908 | 966.7969 | 966.7969 |
| R1_42_1 | 975 | 0.23605 | 0.23605 | R2_59_2 | 2484 | 2.190399 | 35.046384 |  |  |  |  |
| R1_42_2 | 604 | 0.23605 | 0.23605 | R2_41_2 | 616 | 226.593 | 3.77655 |  |  |  |  |
| R2_153_2 | 922 | 0.273789 | 0.136909 | R2_154_1 | 912 | 241.699225 | 241.699225 |  |  |  |  |
| R1_83_2 | 945 | 0.311553 | 0.311553 | R2_7_1 | 635 | 241.699225 | 45.3186 |  |  |  |  |
| R1_81_1 | 889 | 0.316307 | 0.155793 | R2_60_2 | 1625 | 2.492523 | 80.969232 |  |  |  |  |
| R1_12_1 | 1310 | 0.4130875 | 0.9441375 | R2_38_4 | 573 | 258.134746 | 258.134746 |  |  |  |  |
| R2_20_2 | 1343 | 0.44259375 | 1.77028125 | R2_72_2 | 1792 | 259.101569 | 259.101569 |  |  |  |  |
| R2_27_1 | 2254 | 0.47206875 | 0.05900625 | R1_23_1 | 1689 | 26.43585 | 26.43585 |  |  |  |  |
| R2_55_3 | 782 | 0.472075 | 0.354075 | R1_71_2 | 476 | 26.43585 | 26.43585 |  |  |  |  |
| R2_143_1 | 825 | 0.4721 | 7.5531 | R2_14_2 | 596 | 280.371101 | 280.371101 |  |  |  |  |
| R2_153_1 | 664 | 0.538137 | 0.269097 | R1_91_2 | 954 | 28.324125 | 7.5531 |  |  |  |  |
| R1_72_1 | 988 | 0.623139 | 0.03894 | R1_62_1 | 955 | 2.83245 | 0.236025 |  |  |  |  |
| R2_151_2 | 1000 | 0.623139 | 0.623139 | R1_41_2 | 1001 | 30.2124 | 181.2744 |  |  |  |  |
| R2_151_3 | 492 | 0.623139 | 0.623139 | R1_22_2 | 1041 | 30.2124 | 28.324125 |  |  |  |  |
| R1_83_1 | 1648 | 0.632547 | 0.632547 | R1_31_2 | 1466 | 30.2124 | 30.2124 |  |  |  |  |
| R2_151_1 | 794 | 0.642022 | 0.642022 | R2_117_1 | 1039 | 30.2124 | 30.2124 |  |  |  |  |
| R1_43_2 | 2907 | 0.8260875 | 0.9441375 | R2_117_3 | 913 | 30.2124 | 30.2124 |  |  |  |  |
| R1_52_2 | 1504 | 0.88509375 | 0.9441375 | R2_38_1 | 829 | 31.9042944 | 31.9042944 |  |  |  |  |
| R2_1_1 | 995 | 0.9441 | 0.9441 | R2_53_2 | 1989 | 35.046384 | 280.371101 |  |  |  |  |
| R2_57_1 | 1392 | 0.9441375 | 0.8260875 | R2_71_1 | 1127 | 3.5405625 | 3.5405625 |  |  |  |  |
| R2_24_2 | 2985 | 0.9441375 | 0.9441375 | R2_6_1 | 546 | 362.5488 | 60.4248 |  |  |  |  |
| R1_62_2 | 627 | 0.94415 | 0.708075 | R1_101_2 | 430 | 3.77655 | 1.416225 |  |  |  |  |
| R2_63_1 | 1571 | 0.94415 | 0.94415 | R1_32_2 | 783 | 3.77655 | 3.5405625 |  |  |  |  |
| R2_63_3 | 712 | 0.94415 | 0.94415 | R1_23_2 | 1182 | 3.77655 | 3.77655 |  |  |  |  |
| R2_45_1 | 1379 | 0.9970092 | 0.9970092 | R1_71_1 | 825 | 3.77655 | 3.77655 |  |  |  |  |
| R2_19_2 | 6943 | 10.121154 | 1.246278 | R2_67_1 | 523 | 3.7766 | 15.1062 |  |  |  |  |

**Table (S10).**
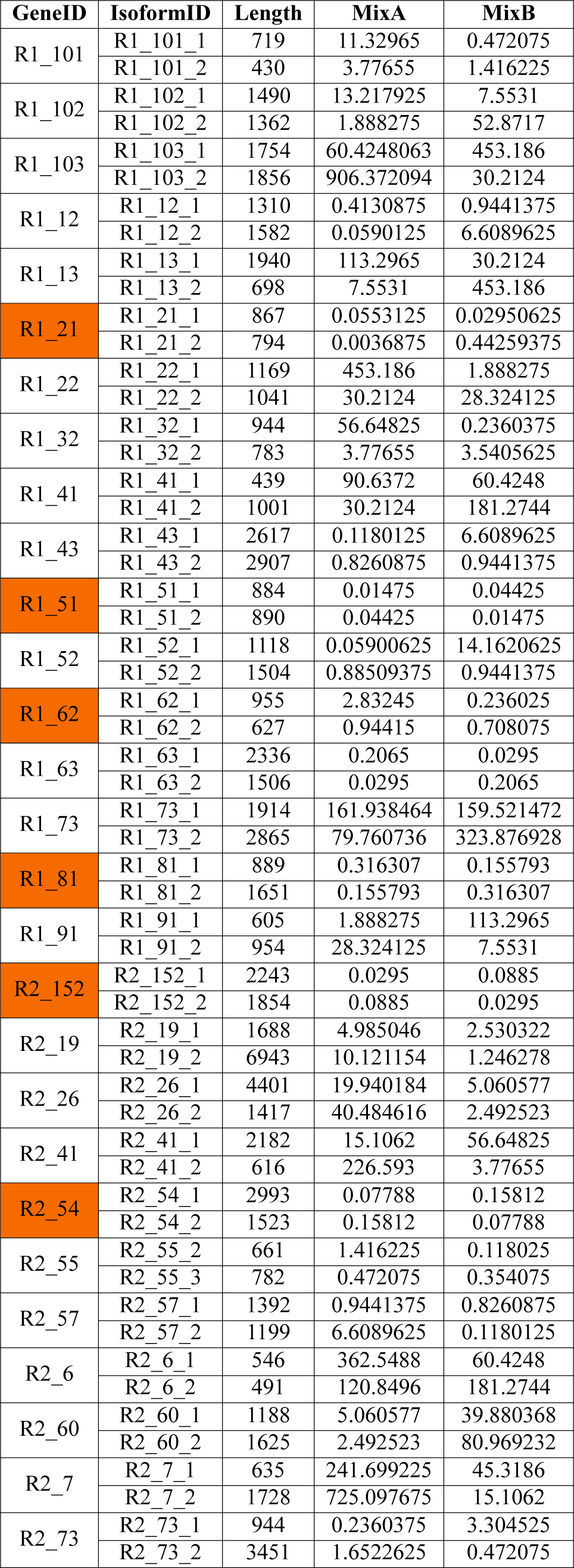
Sequins for differential alternative splicing events. In the table true positive sequins for alternative splicing (AS) events are shown. The two mixtures (in attomoles/*µ*l units) with their respective concentrations are shown. Overall there are 28 true positive sequins for AS events. The saffron colored sequins are true positive, however these were consistently found to be of low coverage regardless of the aligner used for alignment (see Appendix Table). Not to penalize AS event detection tools, the true positive sequins were reset to 22 counts. Sequins with one isoform or no change in ratios across the two mixtures were our true negative. We don’t expect these sequins to be reported by any AS tools. Appendix Figure S3 and Figure S4 show the read coverage from two aligners (STAR and HISAT2).

**Table (S11).** Sequins missed by the DTE tools. The expected absolute log-fold-change values of each sequins is shown in the table.

| Sequins | Total_Isoform | Kallisto | EBSeq | LFC | Tissue |
| --- | --- | --- | --- | --- | --- |
| R1_51 | 2 | Not Reported | Not Reported | 1.5 | Brain, Kidney, Muscle, Retina, RPE |
| R1_81 | 2 | Not Reported | NA | 1.02 | Brain |
| R1_61 | 2 | Not Reported | NA | 0.4 | Muscle |
| R1_21 | 2 | Not Reported | Not Reported | 6.90 | Retina |
| R2_115 | 2 | Not Reported | Not Reported | 1 | RPE |
| R2_54 | 2 | Not Reported | Not Reported | 1.02 | RPE |

**Figure (S3).**
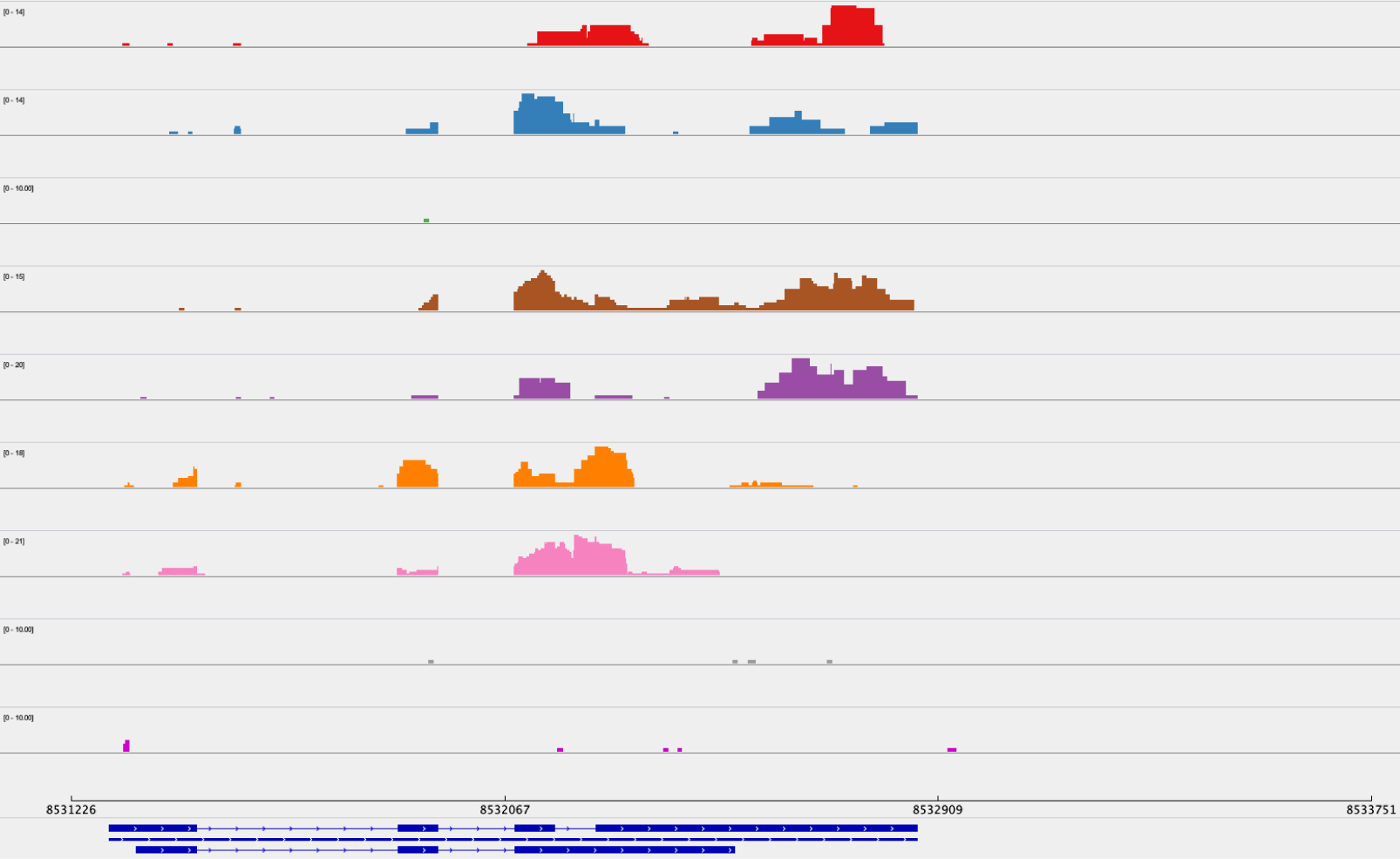
Sashimi plot for true positive sequin using STAR on mouse brain samples. This sequin, R1_62 is not among the lowest concentration of sequin (8th highest concentration) gene. This sequin is not well covered by the aligner.

**Figure (S4).**
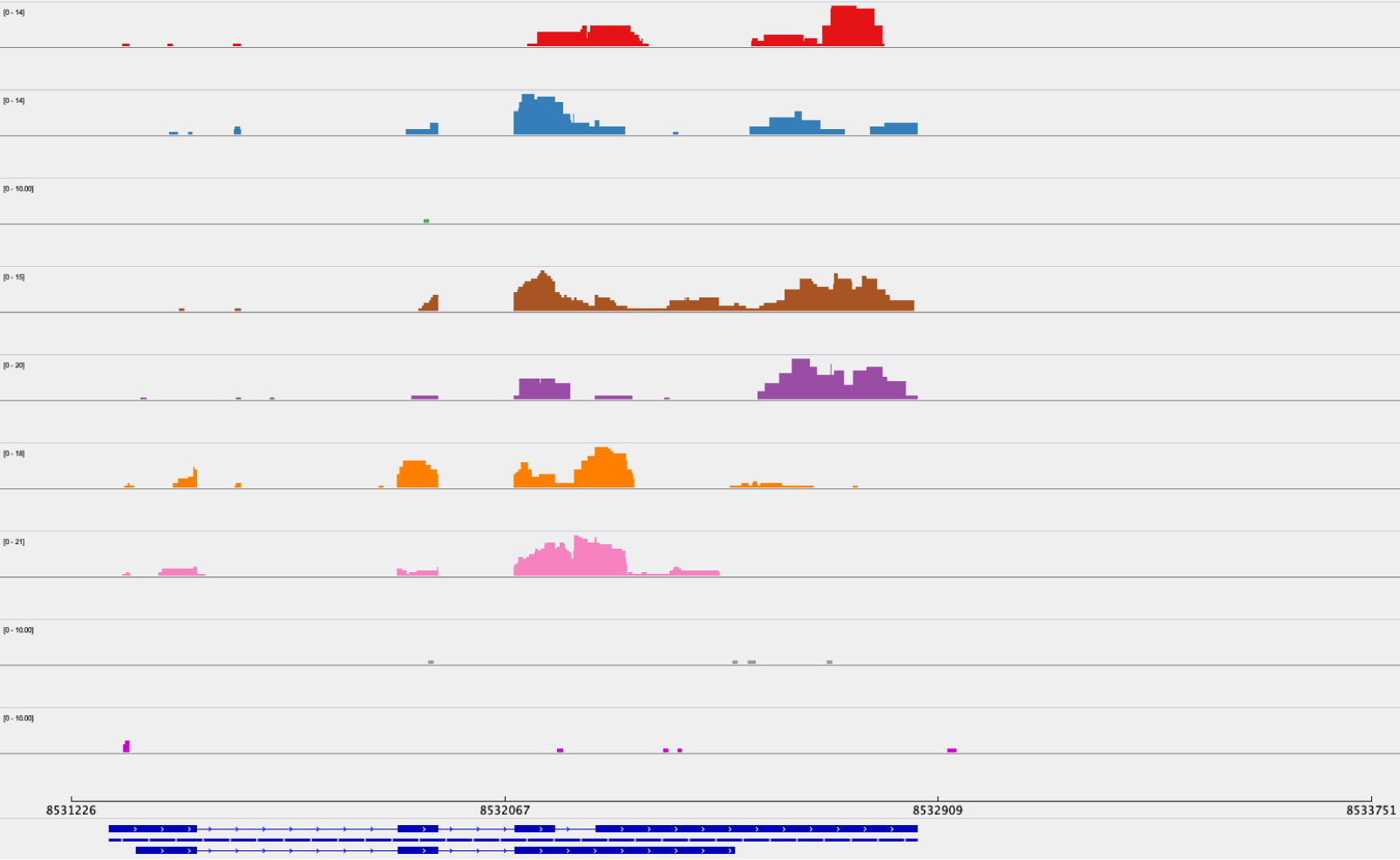
Sashimi plot for true positive sequin using HISAT2 on mouse brain samples. This sequin, R1_62 is not among the lowest concentration of sequin (8th highest concentration) gene. This sequin is not well covered by the aligner.

**Figure (S5).**
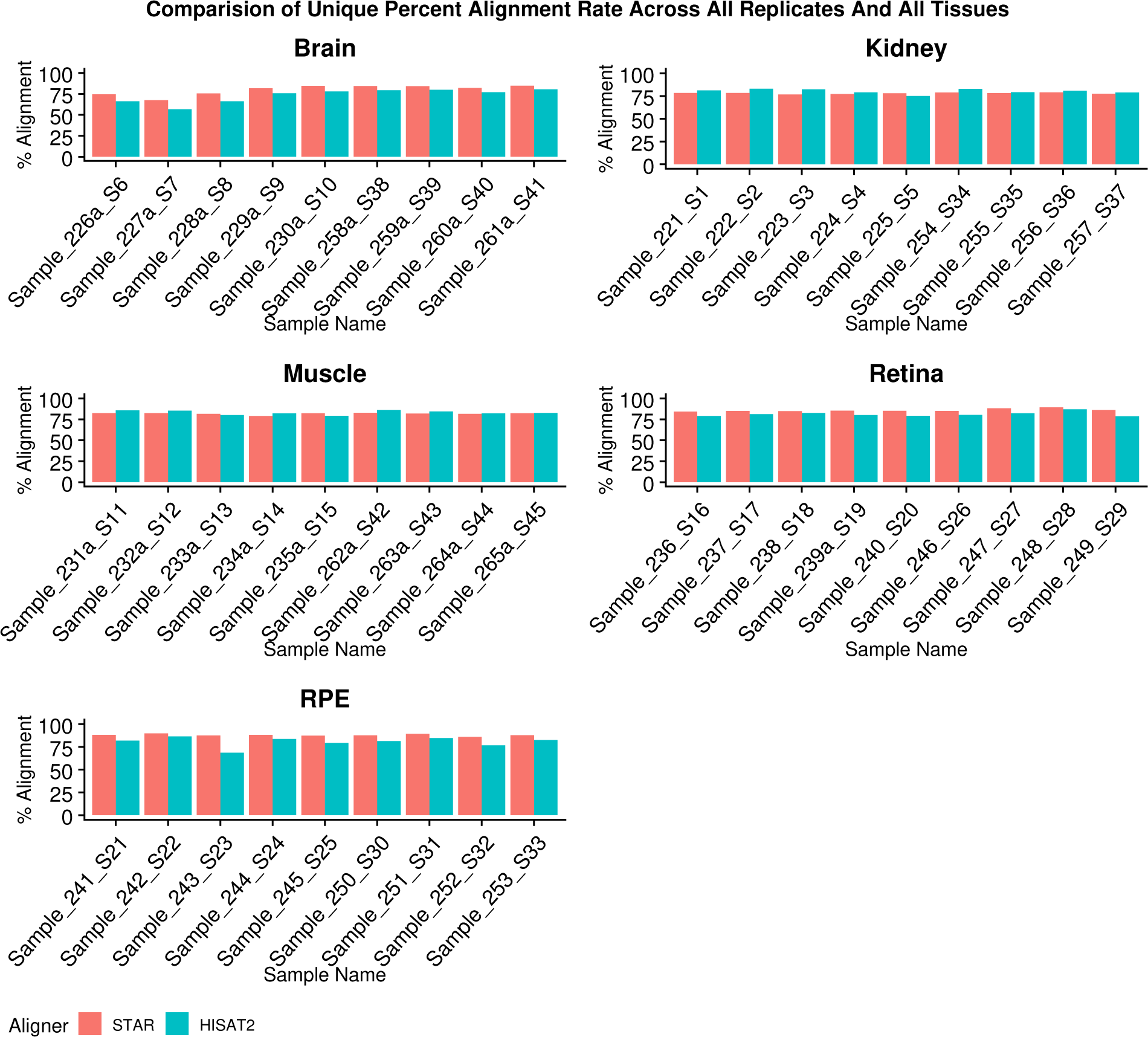
Comparative alignment rate from the two aligners for all the reads. The figure shows percentage unique alignment as reported by STAR and HISAT2 for all the five mouse tissues. Over all the alignment rates were indistinguishable. Assigned unique sample identifier assigned to each sample is shown in the x-axis and percentage alignment rate on y-axis. From left to right, first 5 are mutants followed by wild type samples.

**Figure (S6).**
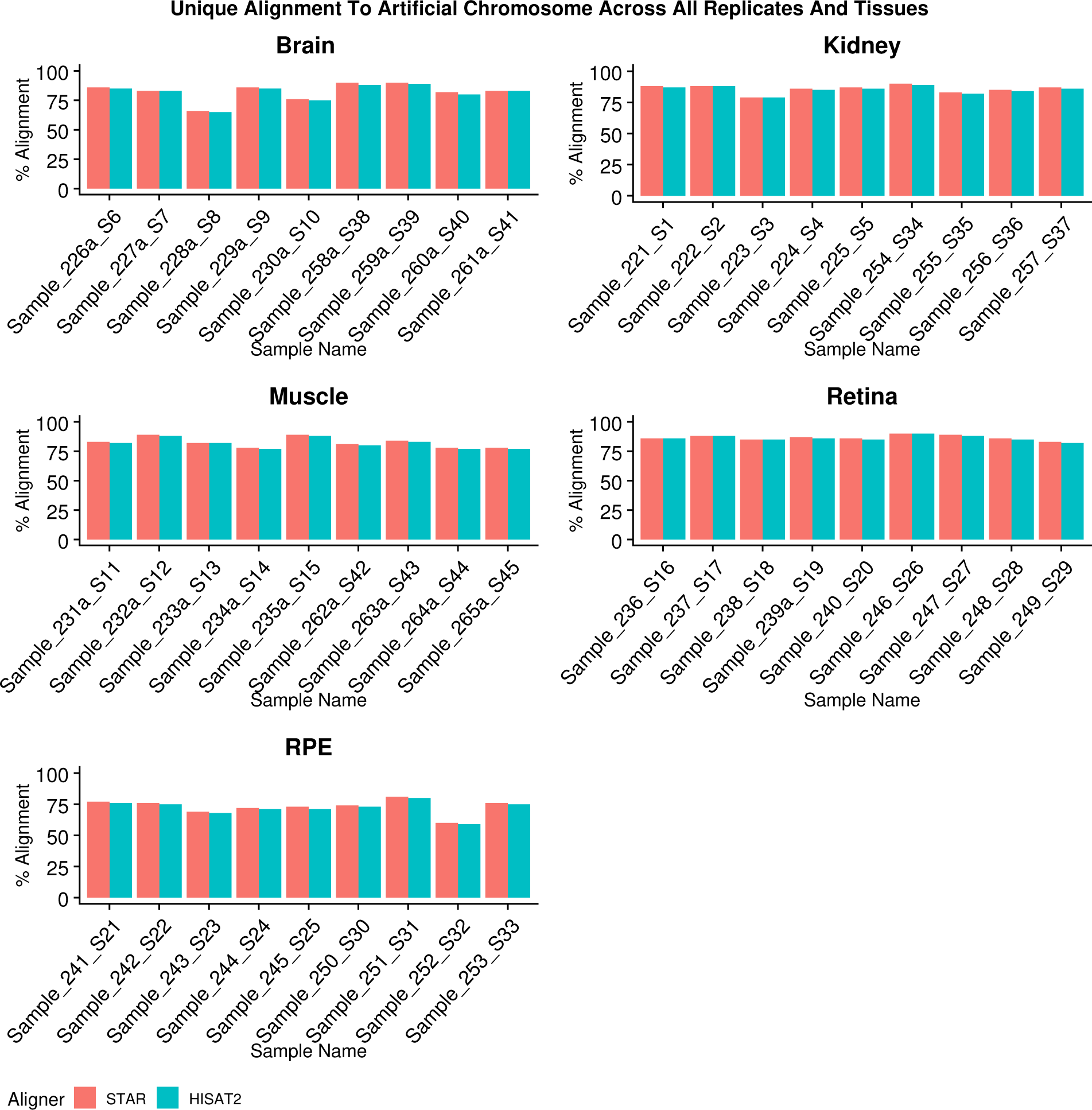
Comparative *IS* specific alignment rate from the two aligners. The figure shows percentage unique alignment specific to the spiked-in *in-silico* artificial chromosome (IS) as reported by STAR and HISAT2 for all the five mouse tissues. The alignment rates were indistinguishable. Assigned unique sample identifier assigned to each sample is shown in the x-axis and percentage alignment rate on y-axis.

**Figure (S7).**
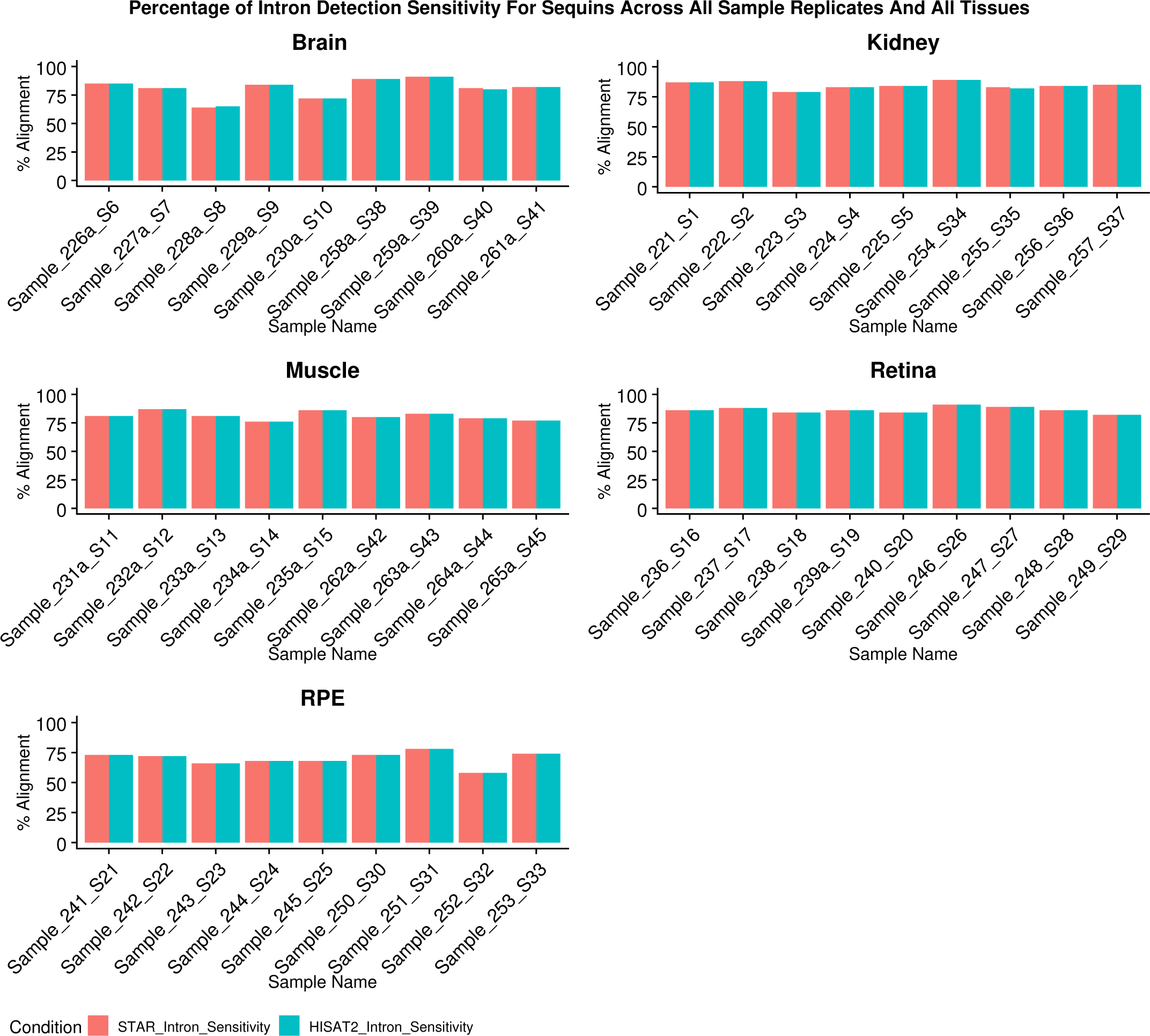
Comparative intron sensitivity for the two aligners. The figure shows intron sensitivity rate as reported by STAR and HISAT2 for all the five mouse tissues using Anaquin. Sensitivity indicates the fraction of annotated regions covered by alignments of the reads. The alignment rates were indistinguishable. From left to right, first 5 are mutants followed by wild type samples.

**Figure (S8).**
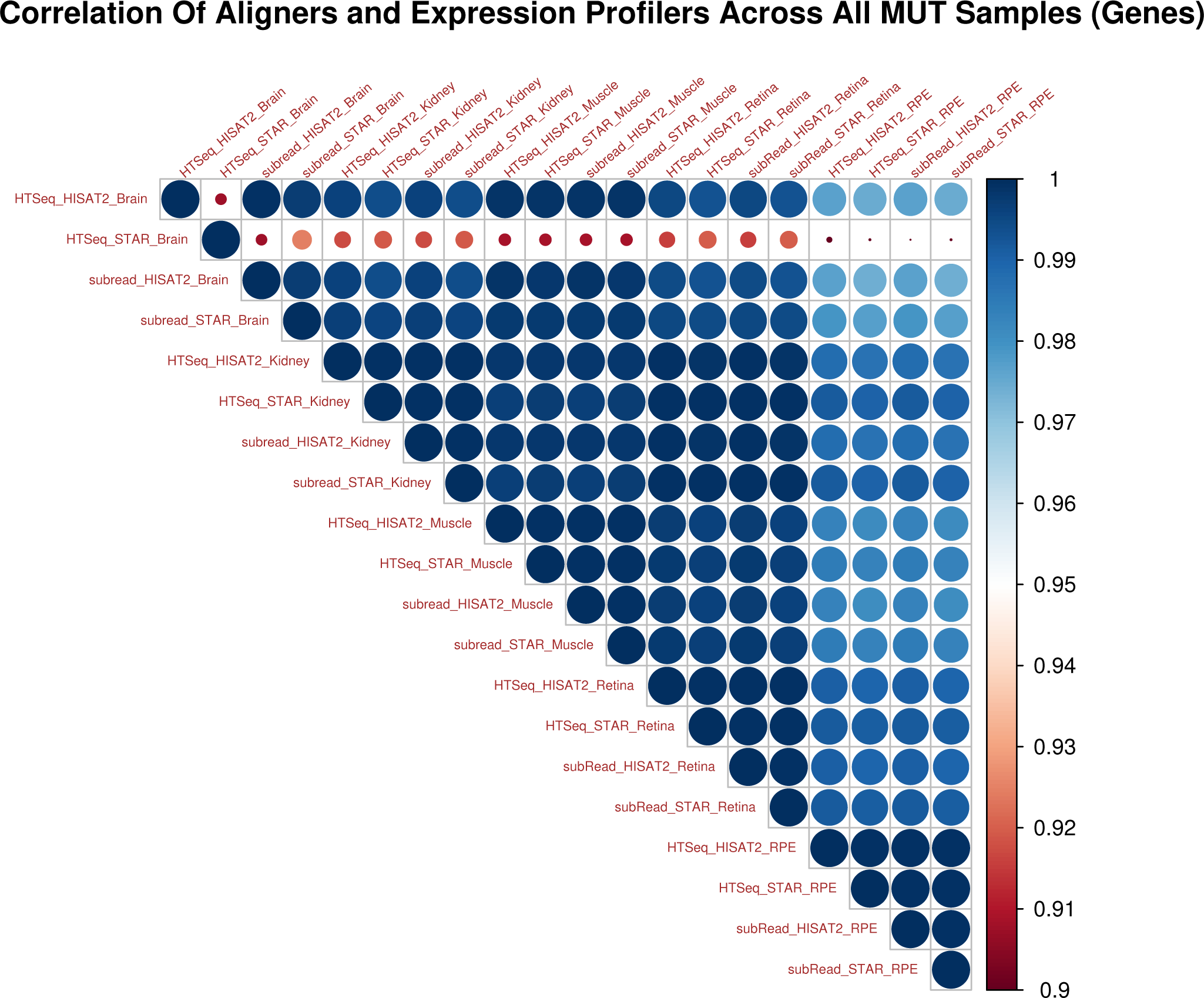
Correlation plot for comparative analysis across all the mutant type tissues. An upper triangular heatmap shows high correlation across different combination of the aligners and read counting tools across the five mouse tissues and their replicates. The average TPM values across the replicates were used to calculate the Pearson correlation.The combinations of tools used are shown in the format of: Gene Expression Profile_Aligner_Tissue.

The following analysis tool combinations were identified (the combination of tools are listed as DGE_aligner_gene expression profile tool, along with missing number of sequin genes in brackets) edgeR_STAR_fC (1), edgeR_STAR_HTC (3), DESeq2_STAR_fC (1) and DESeq2_STAR_HTC (3). Here, fC refers to featureCount and HTC refers to HTSeq-Counts. Similarly, for the kidney sample, the same combinations of analysis tools; edgeR_STAR_fC (1), edgeR_STAR_HTC(1), DESeq2_STAR_fC (1) and DESeq2_STAR_HTC (1) showed drop in performance. The same unary sequin gene, identified in the brain samples, was also not reported in the kidney samples.

**Figure (S9).**
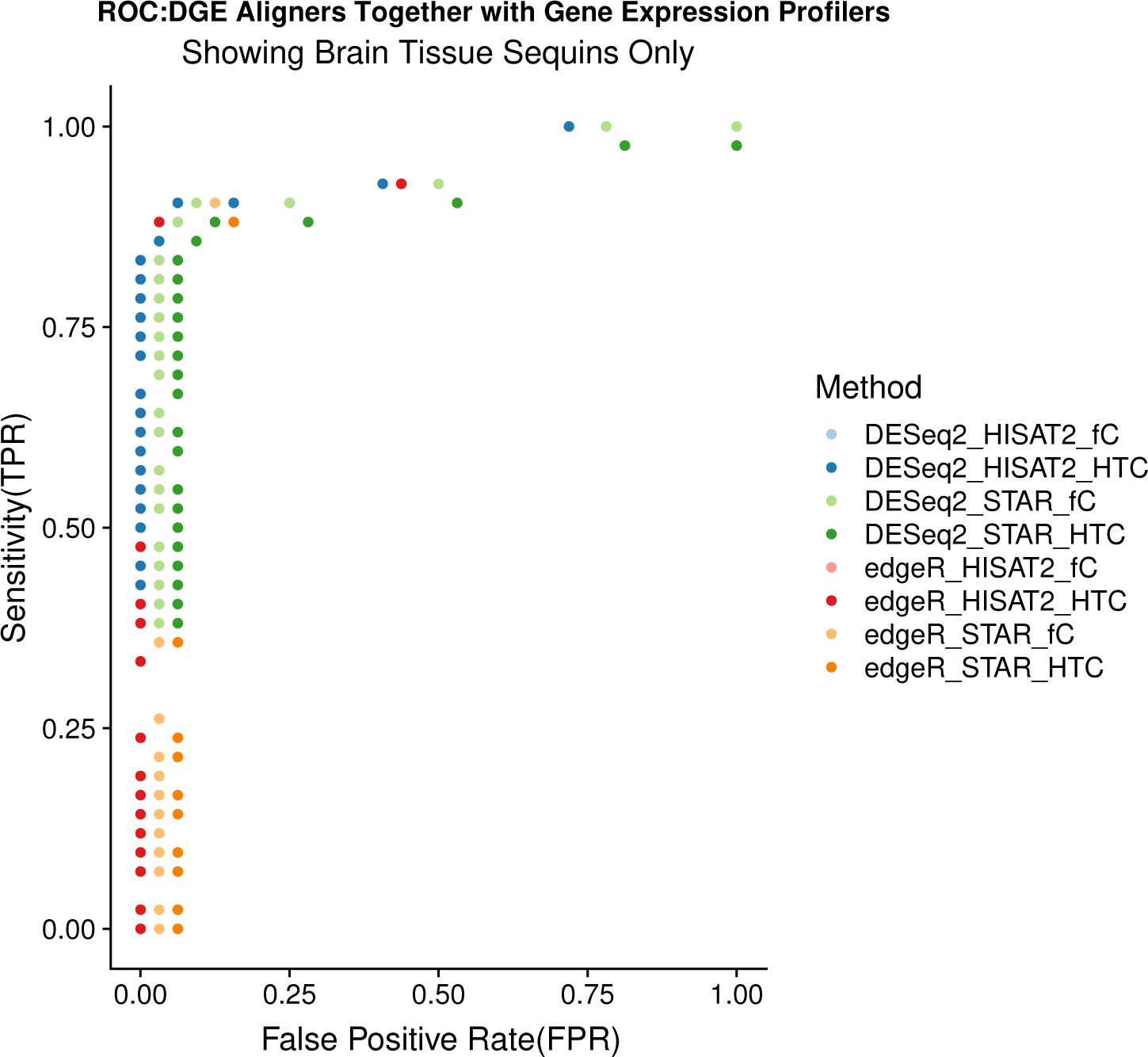
ROC (receiver operating characteristic) curves for Sequin genes for diagnostic performance in mouse brain tissue. The following tools, presented in the form of is DGE_aligner_gene expression profile tool showed poor performance (in brackets total count of missed sequin gene is shown): edgeR_STAR_fC (1), edgeR_STAR_HTC (3), DESeq2_STAR_fC (1) and DESeq2_STAR_HTC (3) compared to rest of the combinations.

**Figure (S10).**
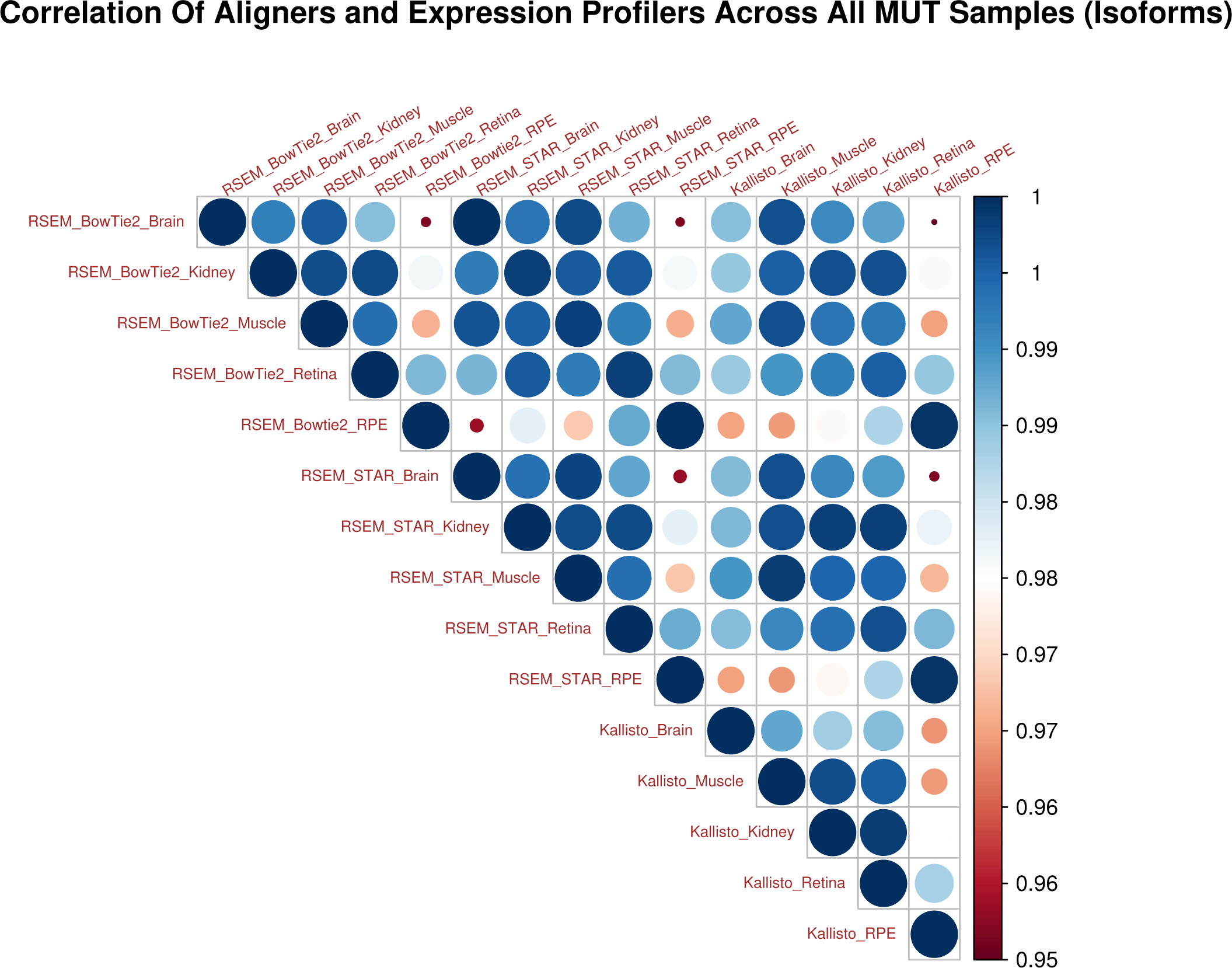
Correlation plot for comparative analysis for all the mutant samples using RSEM and two aligners along with Kallisto across all the mouse tissues. An upper triangular heatmap is generated showing Pearson correlation of determination values using average TPM expression values across the five mouse tissues. The combinations of tools used are shown in the format of: ExpressionTool_Aligner_Tissue, for example, RSEM_STAR_Brain. High correlation of determination values were observed across different tissues types and combinations of different tools.

**Figure (S11).**
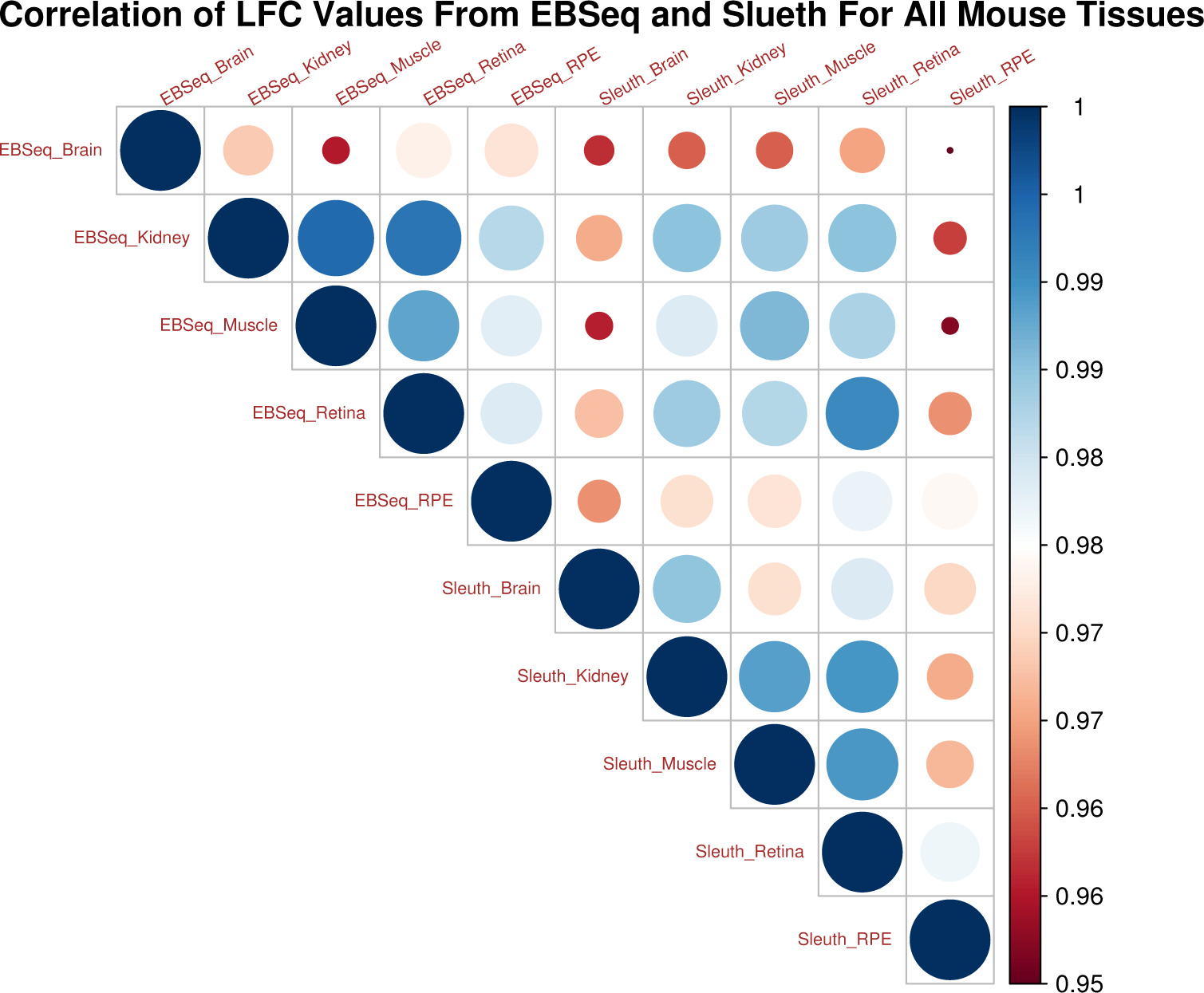
Correlation for differential isoform expression across all five tissues using EBSeq and Kallisto. High correlation was observed for sequins reported by both the tools. We used a “complete case analysis” approach and removed sequins not reported by either of the programs. Refer to Figure S12 for impacts of directionality and proportions and Figure 10 shows sensitivity and specificity including impacts due to missed sequin isoforms.

**Figure (S12).**
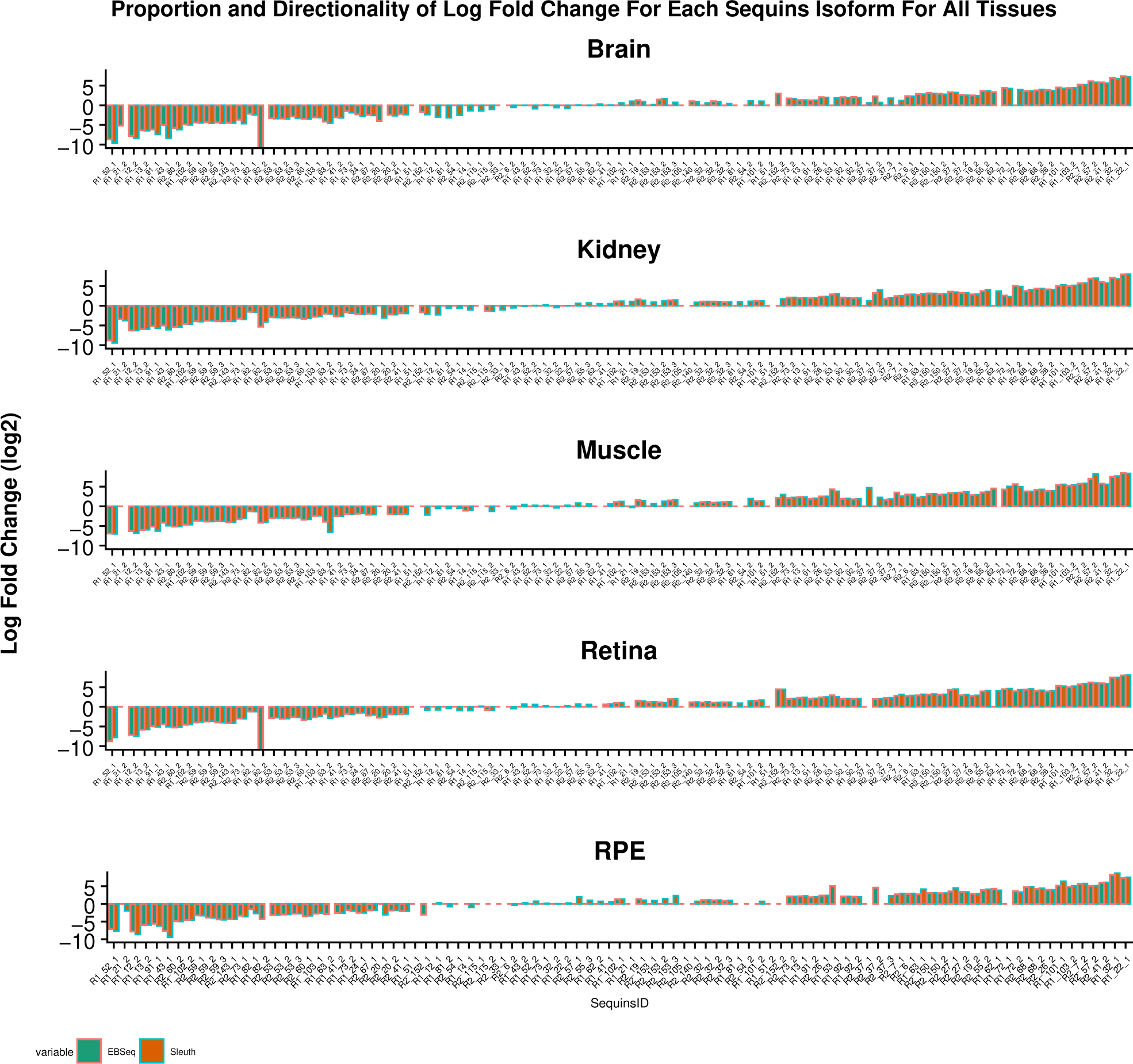
Comparative proportion of log-fold-changes for the sequin genes across all the mouse tissues using different tools. RSEM with default aligner(Bowtie2) was used in conjunction with EBSeq for differential isoform expression. The combined pipeline of Kallisto and Sleuth was used. In the x-axis all the sequin genes are shown and on the y-axis LFC values (log2) are shown. The sequin identifiers are sorted based on expected LFC values and are static all across the tissues. For all mouse tissues, Sleuth and EBSeq missed to report both the isoforms for one sequins. Overall in the brain 4 sequin isoforms, 2 in kidney, 4 in muscle and retina and 6 in RPE were not reported by Kallisto. Similarly, 2 isoforms in the retina and 4 in RPE were not reported by EBSeq. See Table S11 for more details.

**Table (S12).** Sequins sensitivity from two aligners. The table shows counts of AS events from STAR and HISAT2. MAJIQ was used for AS detection. In the mouse retina and brain the percent sensitivity from HISAT2 was slightly higher than STAR. Absence of unary sequins from STAR made the difference in favor of HISAT. In all the other cases it was identical. In caption H refers to HISAT2 and S refers to STAR aligner.

|  | RPE_H | RPE_S | Retina_H | Retina_S | Brain_H | Brain_S | Muscle_H | Muscle_S | Kidney_H | Kidney_S |
| --- | --- | --- | --- | --- | --- | --- | --- | --- | --- | --- |
| **TP** | 15 | 15 | 20 | 19 | 14 | 13 | 17 | 17 | 20 | 20 |
| **FN** | 7 | 7 | 2 | 3 | 8 | 9 | 5 | 5 | 2 | 2 |
| **Sensitivity%** | 68.18 | 68.18 | 90.90 | 86.36 | 63.63 | 59.09 | 77.27 | 77.27 | 90.90 | 90.90 |

**Table (S13).** Sequins sensitivity and specificity from two aligners with JunctionSeq. The table shows counts of AS events from STAR and HISAT2. JunctionSeq was used for AS detection. With exception of reporting of higher false positives for retina samples (5), and muscle (2) near identical results were observed between the two aligners. In caption H refers to HISAT2 and S refers to STAR aligner.

|  | RPE_H | RPE_S | Retina_H | Retina_S | Brain_H | Brain_S | Muscle_H | Muscle_S | Kidney_H | Kidney_S |
| --- | --- | --- | --- | --- | --- | --- | --- | --- | --- | --- |
| **TP** | 20 | 19 | 22 | 22 | 22 | 22 | 22 | 22 | 22 | 22 |
| **FN** | 2 | 3 | 0 | 0 | 0 | 0 | 0 | 0 | 0 | 0 |
| **Sensitivity%** | 90.90 | 86.3 | 100 | 100 | 100 | 100 | 100 | 100 | 100 | 100 |
| **TN** | 46 | 46 | 42 | 46 | 46 | 46 | 45 | 47 | 46 | 47 |
| **FP** | 1 | 1 | 5 | 1 | 1 | 1 | 2 | 0 | 1 | 0 |
| **Specificity%** | 97.87 | 97.87 | 89.36 | 97.87 | 97.87 | 97.87 | 95.74 | 100 | 97.87 | 100 |

**Table (S14).** Novel splice junctions on sequins by two aligners with JunctionSeq. Novel splice junctions on sequins were observed across all the tissues by both the aligners. We noticed reports of novel splice junctions on sequins, more from the STAR aligner. Three (3) was the max number of novel splice junction per gene. SJ refers to splice junctions. Use of STAR aligner with the tissue is marked and unmarked is with HISAT2

| Tissue/Junctions | RPE STAR | RPE | Retina STAR | Retina | Brain STAR | Brain | Kidney STAR | Kidney | Muscle STAR | Muscle |
| --- | --- | --- | --- | --- | --- | --- | --- | --- | --- | --- |
| **Total_Novel_SJ** | 1 | 0 | 6 | 3 | 4 | 1 | 6 | 3 | 5 | 1 |

**Table (S15).**
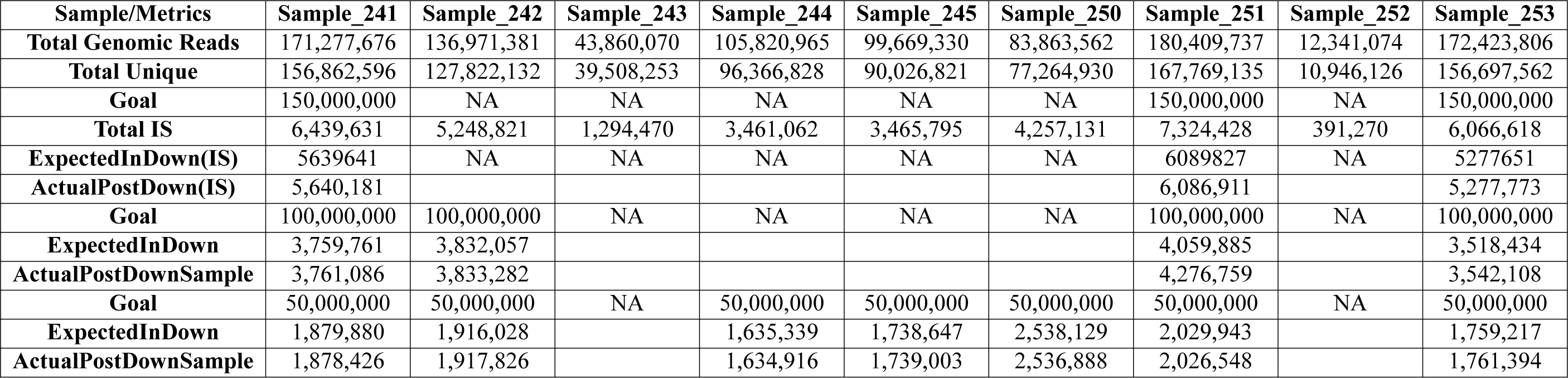
Metrics associated with downsampling mouse RPE sample set. The table shows a number of metrics: original genomic read counts, unique read counts, sequins (IS) read counts, expected IS reads and actual IS reads pre and post downsampling. There were samples that do not meet the minimum read count requirement, such samples were used entirely and thus “NA” is marked where appropriate.

**Table (S16).** Metrics associated with downsampling mouse retina sample set. The table shows a number of metrics: original genomic read counts, unique read counts, sequins (IS) read counts, expected IS reads and actual IS reads pre and post downsampling. There were samples that do not meet the minimum read count requirement, such samples were used entirely and thus “NA” is marked where appropriate

| Sample/Metrics | Sample_236 | Sample_237 | Sample_238 | Sample_239a | Sample_240 | Sample_246 | Sample_247 | Sample_248 | Sample_249 |
| --- | --- | --- | --- | --- | --- | --- | --- | --- | --- |
| **Total Genomic Reads** | 179,710,867 | 202,798,068 | 139,679,922 | 172,407,601 | 137,407,045 | 239,497,635 | 203,542,171 | 188,923,358 | 150,278,217 |
| **Total UniqueReads** | 156,960,473 | 178,562,546 | 124,283,827 | 152,415,475 | 123,087,638 | 214,946,863 | 188,611,822 | 177,222,027 | 134,343,354 |
| **Goal** | 150,000,000 | 150,000,000 | NA | 150,000,000 | NA | 150,000,000 | 150,000,000 | 150,000,000 | NA |
| **Total IS** | 6,503,186 | 7,544,866 | 6,868,886 | 6,450,277 | 7,167,630 | 13,673,884 | 10,238,750 | 9,498,976 | 5,496,948 |
| **ExpectedInDown(IS)** | 5,428,041 | 5,580,575 | NA | 5,611,943 | NA | 8,564,104 | 7,545,427 | 7,541,928 | NA |
| **ActualPostDown(IS)** | 5,427,538 | 5,581,853 |  | 5,611,494 |  | 8,564,204 | 7,546,849 | 7,542,031 |  |
| **Goal** | 100,000,000 | 100,000,000 | 100,000,000 | 100,000,000 | 100,000,000 | 100,000,000 | 100,000,000 | 100,000,000 | 100,000,000 |
| **ExpectedInDown(IS)** | 3618693 | 3720383 | 4917590 | 3741295 | 5216348 | 5709402 | 5030284 | 5027952 | 3657847 |
| **ActualPostDown(IS)** | 3,618,070 | 3,719,887 | 4,919,517 | 3,738,769 | 5,219,733 | 5,711,413 | 5,029,642 | 5,027,127 | 3,656,115 |
| **Goal** | 50,000,000 | 50,000,000 | 50,000,000 | 50,000,000 | 50,000,000 | 50,000,000 | 50,000,000 | 50,000,000 | 50,000,000 |
| **ExpectedInDown(IS)** | 1809346 | 1860191 | 2458795 | 1870647 | 2608174 | 2854701 | 2515142 | 2513976 | 1828923 |
| **ActualPostDown(IS)** | 1,808,600 | 1,860,175 | 2,459,651 | 1,869,461 | 2,605,755 | 2,854,939 | 2,515,147 | 2,516,084 | 1,826,335 |

**Table (S17).** Metrics associated with downsampling mouse brain sample set. The table shows a number of metrics: original genomic read counts, unique read counts, sequins (IS) read counts, expected IS reads and actual IS reads pre and post downsampling. There were samples that do not meet the minimum read count requirement, such samples were used entirely and thus “NA” is marked where appropriate.

| Sample/Metrics | Sample_226a | Sample_227a | Sample_228a | Sample_229a | Sample_230a | Sample_258a | Sample_259a | Sample_260a | Sample_261a |
| --- | --- | --- | --- | --- | --- | --- | --- | --- | --- |
| **Total Genomic Reads** | 224,175,418 | 298,289,266 | 254,811,577 | 167,382,931 | 143,008,334 | 228,263,098 | 193,227,601 | 212,318,905 | 188,721,455 |
| **Total Unique** | 172,297,176 | 206,280,820 | 195,104,137 | 142,045,086 | 124,327,978 | 205,547,310 | 175,360,125 | 181,166,892 | 165,768,006 |
| **Goal** | 150,000,000 | 150,000,000 | 150,000,000 | 150,000,000 | NA | 150,000,000 | 150,000,000 | 150,000,000 | 150,000,000 |
| **Total IS** | 7,395,267 | 7,727,831 | 3,239,520 | 6,757,306 | 3,998,469 | 15,226,593 | 15,045,437 | 8,656,728 | 7,005,287 |
| **ExpectedInDown(IS)** | 4,948,313 | 3,886,076 | 1,907,009 | 6,055,551 | NA | 10,005,949 | 11,679,571 | 6,115,844 | 5,567,958 |
| **ActualPostDown(IS)** | 4,947,822 | 3,883,978 | 1,907,141 | 6,056,207 |  | 10,008,758 | 11,673,649 | 6,115,210 | 5,568,327 |
| **Goal** | 100,000,000 | 100,000,000 | 100,000,000 | 100,000,000 | 100,000,000 | 100,000,000 | 100,000,000 | 100,000,000 | 100,000,000 |
| **ExpectedInDown(IS)** | 3,298,875 | 2,590,717 | 1,271,339 | 4,037,034 | 2,795,969 | 6,670,633 | 7,786,381 | 4,077,229 | 3,711,972 |
| **ActualPostDown(IS)** | 3,299,817 | 2,591,329 | 1,271,553 | 4,037,572 | 2,795,638 | 6,669,839 | 7,783,162 | 4,076,058 | 3,709,669 |
| **Goal** | 50,000,000 | 50,000,000 | 50,000,000 | 50,000,000 | 50,000,000 | 50,000,000 | 50,000,000 | 50,000,000 | 50,000,000 |
| **ExpectedInDown(IS)** | 1,649,438 | 1,295,359 | 635,670 | 2,018,517 | 1,397,985 | 3,335,316 | 3,893,190 | 2,038,615 | 1,855,986 |
| **ActualPostDown(IS)** | 1,648,941 | 1,297,001 | 637,197 | 2,018,611 | 1,396,598 | 3,335,296 | 3,897,183 | 2,039,186 | 1,855,929 |

**Figure (S13).**
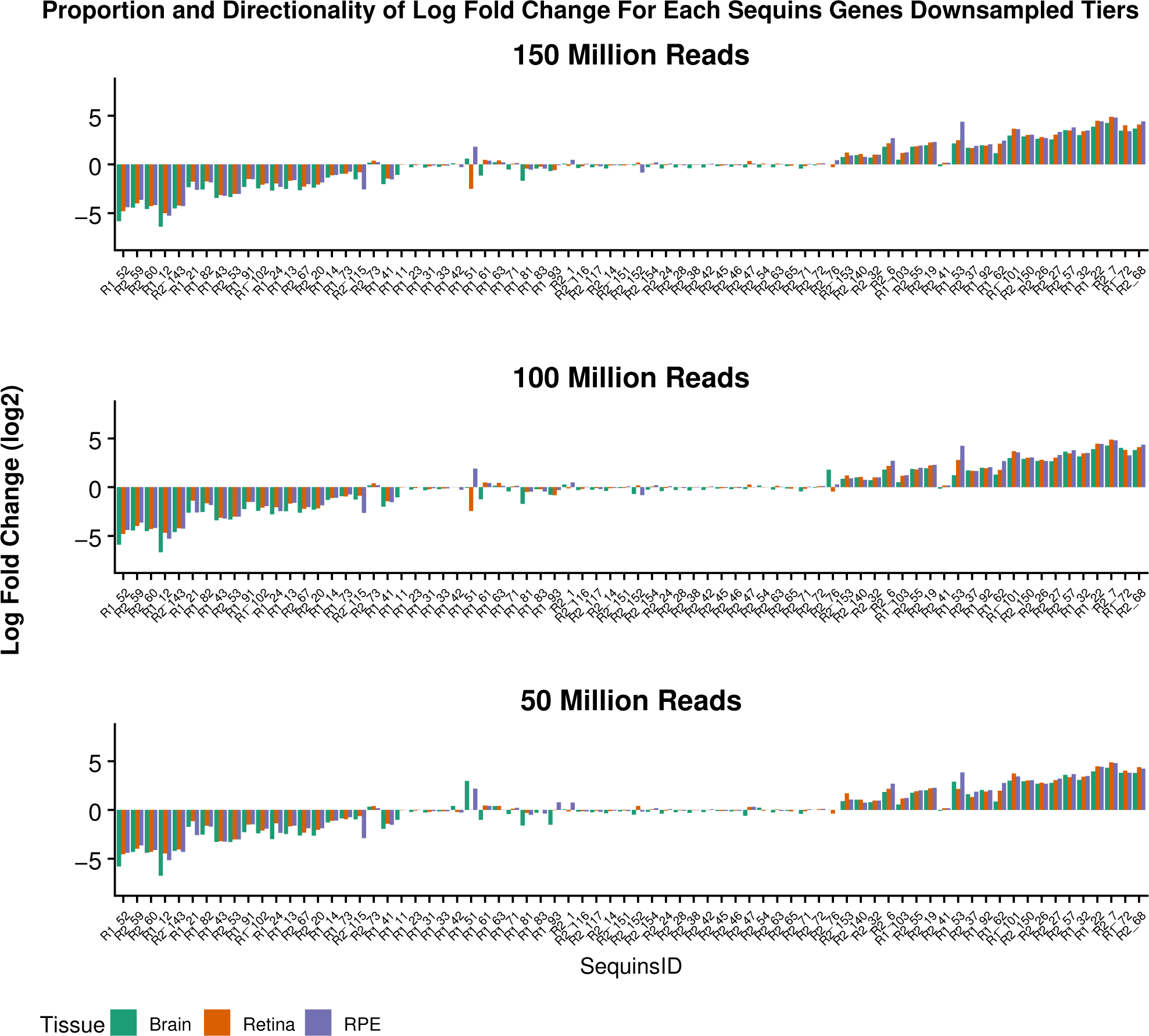
Proportionality and directionality of the downsampled data. The x-axis shows all the sequin genes and y-axis represents the observed LFC values. The observed LFC and directionality for sequin genes across all the three titers is shown. Comparable results were observed regardless of the random read set. As observed with complete read set, sequin genes between −1–+1 showed more variations than the other LFC values. A maximum of 1 sequins were lost. For example, in the 150 million read set, R2_76 in brain and R2_54 in RPE were not reported. In the 100 million read set, R2_54 was not reported in the RPE. In the 50 million read set, R1_51 in retina, R2_54 in RPE and R2_76 in brain and RPE were not reported.

## References

1. Valerio Costa, Marianna Aprile, Roberta Esposito, and Alfredo Ciccodicola. Rna-seq and human complex diseases: recent accomplishments and future perspectives. European Journal of Human Genetics, 21(2):134–142, 2013.

2. Chiara Di Resta, Silvia Galbiati, Paola Carrera, and Maurizio Ferrari. Next-generation sequencing approach for the diagnosis of human diseases: open challenges and new opportunities. Ejifcc, 29(1):4, 2018.

3. Amelia Casamassimi, Antonio Federico, Monica Rienzo, Sabrina Esposito, and Alfredo Ciccodicola. Transcriptome profiling in human diseases: new advances and perspectives. International journal of molecular sciences, 18(8):1652, 2017.

4. Isabelle Audo, Kinga M Bujakowska, Thierry Léveillard, Saddek Mohand-Saïd, Marie-Elise Lancelot, Aurore Germain, Aline Antonio, Christelle Michiels, Jean-Paul Saraiva, Mélanie Letexier, et al. Development and application of a next-generation-sequencing (ngs) approach to detect known and novel gene defects underlying retinal diseases. Orphanet journal of rare diseases, 7(1):8, 2012.

5. Kasper Lage, Niclas Tue Hansen, E Olof Karlberg, Aron C Eklund, Francisco S Roque, Patricia K Donahoe, Zoltan Szallasi, Thomas Skøt Jensen, and Søren Brunak. A large-scale analysis of tissue-specific pathology and gene expression of human disease genes and complexes. Proceedings of the National Academy of Sciences, 105(52):20870–20875, 2008.

6. Dirk A Kleinjan and Veronica van Heyningen. Long-range control of gene expression: emerging mechanisms and disruption in disease. The American Journal of Human Genetics, 76(1):8–32, 2005.

7. Juliana Costa-Silva, Douglas Domingues, and Fabricio Martins Lopes. Rna-seq differential expression analysis: An extended review and a software tool. PloS one, 12(12):e0190152, 2017.

8. Amanda J Ward and Thomas A Cooper. The pathobiology of splicing. The Journal of Pathology: A Journal of the Pathological Society of Great Britain and Ireland, 220(2):152–163, 2010.

9. Nicole Weisschuh, Elena Buena-Atienza, and Bernd Wissinger. Splicing mutations in inherited retinal diseases. Progress in retinal and eye research, page 100874, 2020.

10. Eddie Park, Zhicheng Pan, Zijun Zhang, Lan Lin, and Yi Xing. The expanding landscape of alternative splicing variation in human populations. The American Journal of Human Genetics, 102(1):11–26, 2018.

11. Mihaela Pertea, Geo M Pertea, Corina M Antonescu, Tsung-Cheng Chang, Joshua T Mendell, and Steven L Salzberg. Stringtie enables improved reconstruction of a transcriptome from rna-seq reads. Nature biotechnology, 33(3):290, 2015.

12. Charlotte Soneson, Michael I Love, and Mark D Robinson. Differential analyses for rna-seq: transcript-level estimates improve gene-level inferences. F1000Research, 4, 2015.

13. Giacomo Baruzzo, Katharina E Hayer, Eun Ji Kim, Barbara Di Camillo, Garret A FitzGerald, and Gregory R Grant. Simulation-based comprehensive benchmarking of rna-seq aligners. Nature methods, 14(2):135, 2017.

14. Sayed Mohammad Ebrahim Sahraeian, Marghoob Mohiyuddin, Robert Sebra, Hagen Tilgner, Pegah T Afshar, Kin Fai Au, Narges Bani Asadi, Mark B Gerstein, Wing Hung Wong, Michael P Snyder, et al. Gaining comprehensive biological insight into the transcriptome by performing a broad-spectrum rna-seq analysis. Nature communications, 8(1):1–15, 2017.

15. Arfa Mehmood, Asta Laiho, Mikko S Venäläinen, Aidan J McGlinchey, Ning Wang, and Laura L Elo. Systematic evaluation of differential splicing tools for rna-seq studies. Briefings in Bioinformatics, 2019.

16. Sara Ballouz, Alexander Dobin, Thomas R Gingeras, and Jesse Gillis. The fractured landscape of rna-seq alignment: the default in our stars. Nucleic acids research, 46(10):5125–5138, 2018.

17. Harold Pimentel, Nicolas L Bray, Suzette Puente, Páll Melsted, and Lior Pachter. Differential analysis of rna-seq incorporating quantification uncertainty. Nature methods, 14(7):687, 2017.

18. Shawn C Baker, Steven R Bauer, Richard P Beyer, James D Brenton, Bud Bromley, John Burrill, Helen Causton, Michael P Conley, Rosalie Elespuru, Michael Fero, et al. The external rna controls consortium: a progress report. Nature methods, 2(10):731, 2005.

19. Chi Zhang, Baohong Zhang, Lih-Ling Lin, and Shanrong Zhao. Evaluation and comparison of computational tools for rna-seq isoform quantification. BMC genomics, 18(1):583, 2017.

20. Zhenqiang Su, Paweł P Łabaj, Sheng Li, Jean Thierry-Mieg, Danielle Thierry-Mieg, Wei Shi, Charles Wang, Gary P Schroth, Robert A Setterquist, John F Thompson, et al. A comprehensive assessment of rna-seq accuracy, reproducibility and information content by the sequencing quality control consortium. Nature biotechnology, 32(9):903–914, 2014.

21. Lichun Jiang, Felix Schlesinger, Carrie A Davis, Yu Zhang, Renhua Li, Marc Salit, Thomas R Gingeras, and Brian Oliver. Synthetic spike-in standards for rna-seq experiments. Genome research, 21(9):1543–1551, 2011.

22. Simon A Hardwick, Wendy Y Chen, Ted Wong, Ira W Deveson, James Blackburn, Stacey B Andersen, Lars K Nielsen, John S Mattick, and Tim R Mercer. Spliced synthetic genes as internal controls in rna sequencing experiments. Nature methods, 13(9):792, 2016.

23. Alexander Dobin, Carrie A Davis, Felix Schlesinger, Jorg Drenkow, Chris Zaleski, Sonali Jha, Philippe Batut, Mark Chaisson, and Thomas R Gingeras. Star: ultrafast universal rna-seq aligner. Bioinformatics, 29(1):15–21, 2013.

24. Daehwan Kim, Ben Langmead, and Steven L Salzberg. Hisat: a fast spliced aligner with low memory requirements. Nature methods, 12(4):357, 2015.

25. Yang Liao, Gordon K Smyth, and Wei Shi. featurecounts: an efficient general purpose program for assigning sequence reads to genomic features. Bioinformatics, 30(7):923–930, 2013.

26. Simon Anders, Paul Theodor Pyl, and Wolfgang Huber. Htseq—a python framework to work with high-throughput sequencing data. Bioinformatics, 31(2):166–169, 2015.

27. Michael I Love, Wolfgang Huber, and Simon Anders. Moderated estimation of fold change and dispersion for rna-seq data with deseq2. Genome biology, 15(12):550, 2014.

28. Mark D Robinson, Davis J McCarthy, and Gordon K Smyth. edger: a bioconductor package for differential expression analysis of digital gene expression data. Bioinformatics, 26(1):139–140, 2010.

29. Bo Li and Colin N Dewey. Rsem: accurate transcript quantification from rna-seq data with or without a reference genome. BMC bioinformatics, 12(1):1, 2011.

30. Nicolas L Bray, Harold Pimentel, Páll Melsted, and Lior Pachter. Near-optimal probabilistic rna-seq quantification. Nature biotechnology, 34(5):525, 2016.

31. Ning Leng, John A Dawson, James A Thomson, Victor Ruotti, Anna I Rissman, Bart MG Smits, Jill D Haag, Michael N Gould, Ron M Stewart, and Christina Kendziorski. Ebseq: an empirical bayes hierarchical model for inference in rna-seq experiments. Bioinformatics, 29(8):1035–1043, 2013.

32. Stephen W Hartley and James C Mullikin. Detection and visualization of differential splicing in rna-seq data with junctionseq. Nucleic acids research, 44(15):e127–e127, 2016.

33. Jorge Vaquero-Garcia, Alejandro Barrera, Matthew R Gazzara, Juan Gonzalez-Vallinas, Nicholas F Lahens, John B Hogenesch, Kristen W Lynch, and Yoseph Barash. A new view of transcriptome complexity and regulation through the lens of local splicing variations. elife, 5:e11752, 2016.

34. Kinga Bujakowska, Cecilia Maubaret, Christina F Chakarova, Naoyuki Tanimoto, Susanne C Beck, Edda Fahl, Marian M Humphries, Paul F Kenna, Evgeny Makarov, Olga Makarova, et al. Study of gene-targeted mouse models of splicing factor gene prpf31 implicated in human autosomal dominant retinitis pigmentosa (rp). Investigative ophthalmology & visual science, 50(12):5927–5933, 2009.

35. Eranga N Vithana, Leen Abu-Safieh, Maxine J Allen, Alisoun Carey, Myrto Papaioannou, Christina Chakarova, Mai Al-Maghtheh, Neil D Ebenezer, Catherine Willis, Anthony T Moore, et al. A human homolog of yeast pre-mrna splicing gene, prp31, underlies autosomal dominant retinitis pigmentosa on chromosome 19q13. 4 (rp11). Molecular cell, 8(2):375–381, 2001.

36. Rosario Fernandez-Godino, Donita L Garland, and Eric A Pierce. Isolation, culture and characterization of primary mouse rpe cells. Nature protocols, 11(7):1206–1218, 2016.

37. Babraham Bioinformatics. Fastqc a quality control tool for high throughput sequence data. Cambridge, UK: *Babraham Institute*, 2011.

38. Philip Ewels, Måns Magnusson, Sverker Lundin, and Max Käller. Multiqc: summarize analysis results for multiple tools and samples in a single report. Bioinformatics, 32(19):3047–3048, 2016.

39. Ben Langmead and Steven L Salzberg. Fast gapped-read alignment with bowtie 2. Nature methods, 9(4):357, 2012.

40. Heng Li, Bob Handsaker, Alec Wysoker, Tim Fennell, Jue Ruan, Nils Homer, Gabor Marth, Goncalo Abecasis, and Richard Durbin. The sequence alignment/map format and samtools. Bioinformatics, 25(16):2078–2079, 2009.

41. Taiyun Wei, Viliam Simko, Michael Levy, Yihui Xie, Yan Jin, and Jeff Zemla. Package ‘corrplot’. Statistician, 56(316):e24, 2017.

42. Shihao Shen, Juw Won Park, Zhi-xiang Lu, Lan Lin, Michael D Henry, Ying Nian Wu, Qing Zhou, and Yi Xing. rmats: robust and flexible detection of differential alternative splicing from replicate rna-seq data. Proceedings of the National Academy of Sciences, 111(51):E5593–E5601, 2014.

43. Michael H Farkas, Deborah S Lew, Maria E Sousa, Kinga Bujakowska, Jonathan Chatagnon, Shomi S Bhattacharya, Eric A Pierce, and Emeline F Nandrot. Mutations in pre-mrna processing factors 3, 8, and 31 cause dysfunction of the retinal pigment epithelium. The American journal of pathology, 184(10):2641–2652, 2014.

44. Ted Wong, Ira W Deveson, Simon A Hardwick, and Tim R Mercer. Anaquin: a software toolkit for the analysis of spike-in controls for next generation sequencing. Bioinformatics, 33(11):1723–1724, 2017.

45. Pierre-Luc Germain, Alessandro Vitriolo, Antonio Adamo, Pasquale Laise, Vivek Das, and Giuseppe Testa. Rnaonthebench: computational and empirical resources for benchmarking rnaseq quantification and differential expression methods. Nucleic acids research, 44(11):5054–5067, 2016.

